# A framework for selectively breeding corals for assisted evolution

**DOI:** 10.1101/2021.02.23.432469

**Authors:** Adriana Humanes, John Bythell, Elizabeth Beauchamp, Mitch Carl, Jamie Craggs, Alasdair Edwards, Yimnang Golbuu, Liam Lachs, Janna Leigh Randle, Helios Martinez, Pawel Palmowski, Faith Paysinger, Eveline van der Steeg, Michael Sweet, Achim Treumann, James Guest

## Abstract

Coral cover on tropical reefs has declined during the last three decades due to the combined effects of climate change, destructive fishing, pollution, and land use change. Drastic reductions in greenhouse gas emissions combined with effective coastal management and conservation strategies are essential to slow this decline. Innovative approaches, such as selective breeding for adaptive traits combined with large-scale sexual propagation, are being developed with the aim of pre-adapting reefs to increased ocean warming. However, there are still major gaps in our understanding of the technical and methodological constraints to producing corals for such restoration interventions. Here we propose a framework for selectively breeding corals and rearing them from eggs to 2.5-year old colonies using the coral *Acropora digitifera* as a model species. We present methods for choosing colonies for selective crossing, enhancing early survivorship in *ex situ* and *in situ* nurseries, and outplanting and monitoring colonies on natal reefs. We used a short-term (7-day) temperature stress assay to select parental colonies based on heat tolerance of excised branches. From six parental colonies, we produced 12 distinct crosses, and compared survivorship and growth of colonies transferred to *in situ* nurseries or outplanted to the reef at different ages. We demonstrate that selectively breeding and rearing coral colonies is technically feasible at small scales and could be upscaled as part of restorative assisted evolution initiatives. Nonetheless, there are still challenges to overcome before selective breeding can be implemented as a viable conservation tool, especially at the post-settlement and outplanting phases. Although interdisciplinary approaches will be needed to overcome many of the challenges identified in this study, selective breeding has the potential to be a viable tool within reef managers’ toolbox to support the persistence of selected reefs in the face of climate change.

## Introduction

The Anthropocene, the era in which humans have become a global geophysical force, is characterized by the degradation of ecosystem structure and function, loss of biodiversity and increased rates of species extinction (Steffen et al., 2007; Ceballos et al., 2015). Unfortunately, many existing conservation practices, that are based on local management, are inadequate in the face of new global scale stressors such as those caused by climate change (Lennon, 2015). Coral reefs are among the ecosystems most impacted by human activities and climate change (Hoegh-Guldberg et al., 2019), leading to more rapid increases in extinction risk for many coral species compared to mammals, birds and amphibians (Bongaarts, 2019). During the last 30 years coral cover worldwide has decreased by an estimated 20% (Hoegh-Guldberg et al., 2019), and four pan-tropical coral bleaching events since 1983 have led to coral declines on hundreds of reefs (Lough et al., 2018). The limitations of localised management were most evident in Australia’s Great Barrier Reef Marine Park, between 2015 and 2017. During that period, catastrophic coral bleaching and mortality driven by high sea temperatures occurred despite local management of coral reef fisheries and water quality (Hughes et al., 2017). The present rate of coral reef degradation highlights the urgent need to develop innovative conservation approaches that can maintain ecosystem services and ecological function despite projected sea warming owing to climate change.

As a result of anthropogenic climate change, the frequency, duration, and intensity of marine heat waves increased more than 20-fold between 1981 to 2017 (Laufkötter et al., 2020). Global mean sea surface temperature is projected to reach 1.5 °C above that in pre-industrial times between 2030 and 2052, suggesting that shallow water corals have ∼10 to 30 years to adapt to this temperature increase (Hoegh-Guldberg et al., 2019). For many coral species this period will be too short for adaptation to happen by natural selection, given the sporadic nature of heatwaves at local scales (Bay et al., 2017). Even if warming can be limited to <1.5 °C, it is highly likely that large areas of coral reefs will be experiencing regular mass bleaching events, threatening 70-90% of reefs by 2050 (Hoegh-Guldberg et al., 2018). In addition to tackling climate change, it is increasingly stated that traditional conservation efforts need to be coupled with restoration to assist recovery from disturbances (Anthony et al. 2015). Innovative solutions for actively assisting coral populations to pre-adapt to climate change via assisted evolution (AE) have been proposed to be included in management strategies for coral reefs (van Oppen et al., 2015). The goal of AE is to deliberately enhance certain traits in selected organisms, increasing their chances of surviving in the face of global change (Jones and Monaco, 2009). Such practices may involve induced acclimatisation, modification of the microbial or the Symbiodiniaceae symbiont communities, and selective breeding for adaptive traits.

Selective breeding (SB) is the process by which humans choose individuals with specific heritable phenotypic traits to breed together and produce offspring. Humans have practised SB for centuries to improve the production and taste of crops and livestock (Denison et al., 2003). More recently, such practices have been used to select for traits that might be beneficial in a changing climate such as drought resistance in plants (Hu and Xiong, 2014). SB can also be used as a conservation method for preserving populations of endangered species, however, there are only a few examples where this has been considered as a management strategy (Jones et al., 2007; Aitken and Bemmels, 2016). In marine invertebrates, SB has been used primarily in mollusc aquaculture to improve their growth (Hollenbeck and Johnston, 2018), protein content (Gjedrem et al., 2012), and disease resistance (Parker et al., 2012), highlighting that this approach can be adapted for calcifying organisms. Selecting for heat tolerance in corals is one of the approaches proposed in assisted evolution initiatives, given the benefits that this trait could aid corals in the face of climate change. There is evidence for heritability of heat tolerance in some coral species at early life history stages (e.g., larvae and 70 day old colonies) (Dixon et al., 2015; Kirk et al., 2018; Quigley et al., 2020), suggesting that selective crosses between colonies with known tolerances could produce offspring with above-average heat resistance. To successfully conduct SB, adult colonies with relatively high heat tolerances need to be identified and used as broodstock. These colonies can come from different populations exposed to contrasting temperatures profiles at a range of spatial scales (Dixon et al., 2015; Liew et al., 2020; McClanahan et al., 2020), or among individuals from within a single population where there is sufficient intrapopulation variability. While it is well established that coral populations experiencing higher mean SSTs or more variable temperatures tend to be more tolerant to heat stress (Howells et al., 2012; Thomas et al., 2018), less is known about the extent of within-population variation in heat tolerance for corals (Bay and Palumbi, 2014; van Oppen et al., 2018). However, if sufficient variability does exist then this approach has the advantage of reducing the likelihood of maladaptation to environmental variables other than temperature (Cotto et al., 2019) and may reduce the risk of inadvertently selecting different genetic variants or sub-species.

For AE methods to be successfully incorporated into coral reef resilience adaptation programs, they will need to be combined with techniques to restore and rehabilitate coral reefs (van Oppen et al., 2017). Some of the techniques associated with AE will rely on successful coral larval propagation (CLP) via sexual reproduction. For the purpose of this paper, we define CLP as the process of producing and rearing corals from eggs through to being colonies that are recruited into the population. We define a recruited colony as one that has been transplanted to the reef, has self-attached (sensu Guest et al., 2011), and contributes to the emergent properties of the population (growth, survivorship, and/or reproduction rates). CLP is an emerging method for producing large numbers of corals for reef rehabilitation and restoration, that overcomes early survivorship bottlenecks via a combination of land (*ex situ*) or ocean (*in situ*) based nurseries for rearing the early life stages. Several advances have been made to improve the practices associated with CLP in recent years. The modes of sexual reproduction for approximately half of extant species of hermatypic scleractinians have been identified (Baird et al., 2009) and the timing of spawning catalogued for >300 Indo-Pacific coral species (Baird et al., 2021). CLP has been successfully executed in different geographical regions with different species under *ex situ* and *in situ* conditions (Omori, 2019; Randall et al., 2020) with sexually propagated colonies that were outplanted to the reef reaching sexual maturity (Nakamura et al., 2011; Baria et al., 2012; Guest et al., 2014; Chamberland et al., 2016). Competent coral larvae have been seeded *en masse* onto natural substrates to enhance recruitment (dela Cruz and Harrison, 2017; Doropoulos et al., 2019) and substrates have been designed to settle coral larvae for nursery rearing and outplantation improving early survivorship (Guest et al., 2014; Chamberland et al., 2017). Despite advances in the practice of CLP, most research has focused on the steps during and post-spawning, with little attention given to the provenance or phenotype of the broodstock colonies (apart from laboratory based studies e.g., Dixon et al., 2015; Liew et al., 2020; Quigley et al., 2020). To the best of our knowledge, there have been no attempts to select parents for adaptive traits, such as heat tolerance, as part of CLP for reef rehabilitation.

Here we present a practical framework for developing, testing and implementing SB that can be adapted for coral reef rehabilitation and AE programs. In this study we combine SB trials with coral sexual propagation. We use *Acropora digitifera* as a model species to select for heat tolerance, however this framework can also be applied to other propagule-producing organisms (Vanderklift et al., 2020), and other adaptive traits like growth rate, disease resistance or wound healing capability (Baums et al., 2019). The proposed framework is structured into six sections: 1) selection of parental colonies with traits of interest based on phenotypic or other functional characteristics (e.g. known genotypic markers), 2) design of crosses for SB, 3) methods for collecting gametes to perform SB with corals, 4) methods of larval rearing and settlement onto substrate units *en masse* to produce coral colonies, 5) rearing of coral colonies (*in situ* or *ex situ*) for later outplanting to natural reef habitats, and 6) outplant of corals to the reef and monitoring of their growth and survivorship. Testing for heritability and potential resource trade-offs are also critical steps in SB (Ortiz et al., 2013; Cunning et al., 2015), and are being carried out as part of our ongoing work, however, the results of these studies will be reported elsewhere.

## Materials and methods

### a) Selection of parental colonies for selective breeding

The reef-building coral *Acropora digitifera*, was used as a model for SB as it is widely distributed and abundant on shallow reefs throughout the Indo-West Pacific. Its digitate morphology facilitates fragment removal for conducting stress assays, and spawning times are well established for many locations (Keith et al., 2016; Baird et al., 2021). All of the work described here was carried out at the Palau International Coral Reef Center (PICRC) in the Republic of Palau located in Western Pacific Ocean (Fig. 1A). The source site for all colonies is a shallow, exposed patch-like reef (Mascherchur, N 07°17’ 29.3’’; E 134°31’ 8.00’’, Fig. 1B), where *A. digitifera* is abundant at depths ranging between 0 and 4 m. In November 2017, 99 visibly healthy adult coral colonies were tagged and mapped along eleven 20-m long fixed transects. From these 99 colonies, 34 were randomly selected to assess their performance during a short-term (7-day) temperature stress assay to select parental colonies for the broodstock.

**Figure 1:**
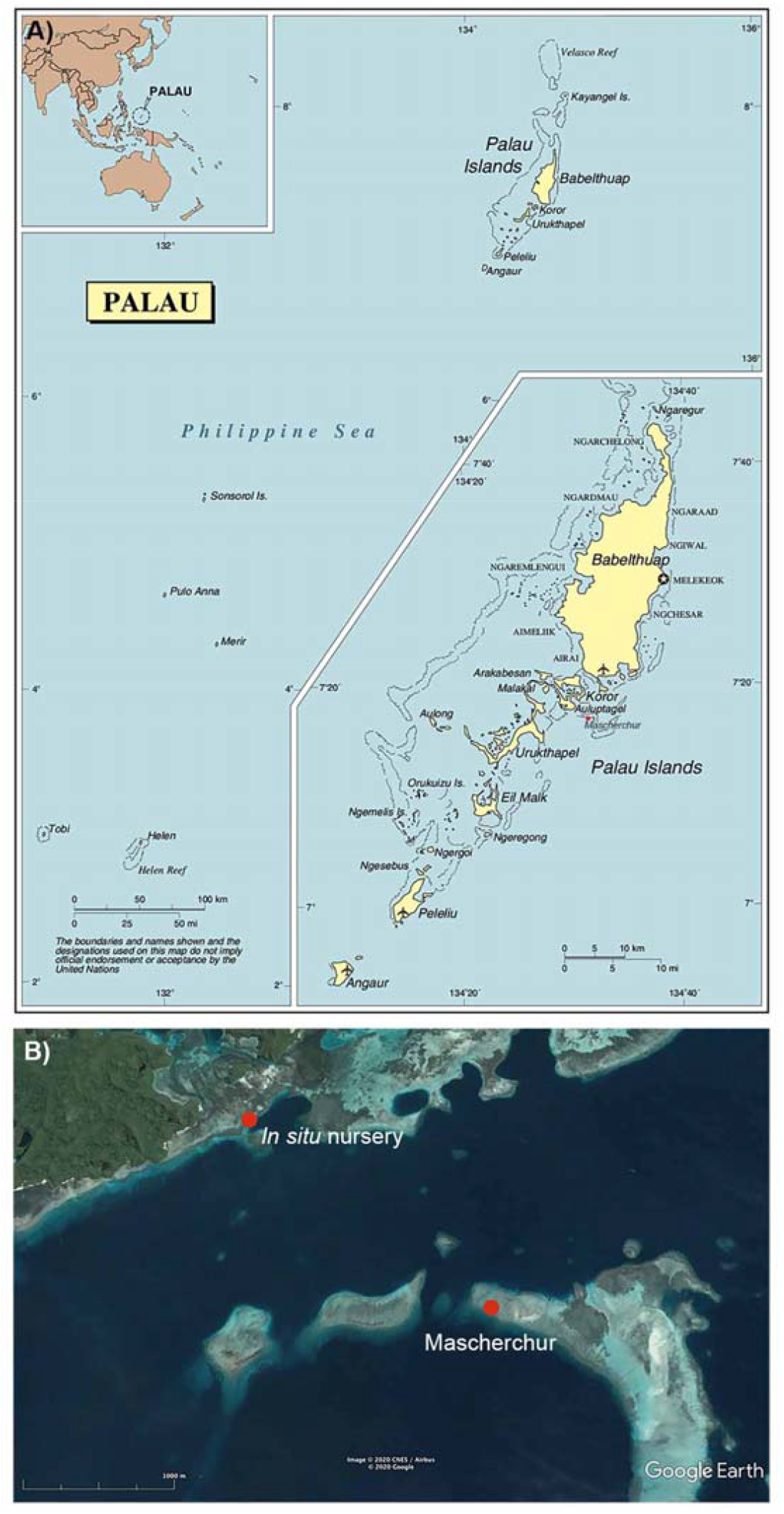
A) Map of Palau modified from https://www.mapsland.com/oceania/palau/large-detailed-political-map-of-palau-with-cities-and-airports. Mascherchur is indicated with a red circle south from Babelthuap Island. B) Map showing the location of the site were transects were deployed at Mascherchur (N 07°17’ 29.3’’; E 134°31’ 8.00’’, indicated with a red circle), and the *in situ* nursery (N 7°18’19.80”N; E 134°30’6.70”E, indicated with a red circle) located 2.2 km away from the study site. Image modified from Google Earth.

For this short-term assay, seven ∼3 cm long fragments were excised from each colony and transported by boat in 50 L seawater tanks to PICRC (∼20 min boat travel time). The donor colonies remained on the reef to recover for approximately five months before SB work began. The 238 fragments were glued to aragonite substrata (∼20 mm diameter, Oceans Wonders LLC) with ethyl cyanoacrylate gel (Coraffix gel), labelled and mounted into plastic holders, that were attached to plexiglass racks (Fig. 2A-J). To determine the relative heat tolerance (RHT) of each colony, a 7-day temperature stress experiment was performed using two temperature levels: a) ambient seawater temperature conditions (30.37 ± 0.46 °C, three replicate tanks, Fig. 2C, F, I), and b) heat stress conditions (Fig. 2A, B, D, E, G, H), where temperature was raised incrementally over the course of three days (+2°C on day one, and +1.5°C on day three), reaching a daily average temperature of 32.95°C (±0.37) during days four to seven (five replicate tanks, SM. 1). Replicate fragments were randomly distributed among seven treatment tanks (24 fragments per tank), with all colonies having at least two replicates in independent thermal stress tanks and at least one replicate in an ambient temperature tank (used as a control for handling stress). The status of each fragment was visually inspected by the same observer daily and ranked as: 1) healthy (no signs of discoloration or mortality), 2) partial mortality (less than 30% of surface area dead) or, 3) dead (more than 30% of the surface with bare skeleton and without tissue). Relative heat tolerance was determined by the end-point mortality (six days after the first temperature increase). Colonies with all replicate stressed fragments alive (0% mortality) were considered to have Relatively High Heat Tolerance (RHHT), whereas colonies with all stressed fragments dead (100% mortality) were classified as having Relatively Low Heat Tolerance (RLHT). Colonies that were not classified either as RHHT or RLHT were considered as unclassified. RHT was considered as unresolved for colonies with control fragments held at ambient temperature that showed a stress response (partial mortality or death), as this might have resulted from handling stress. Relevant National and State permits were obtained for the collection of fragments (National Marine Research Permits: RE-018, RE-18-13).

**Figure 2:**
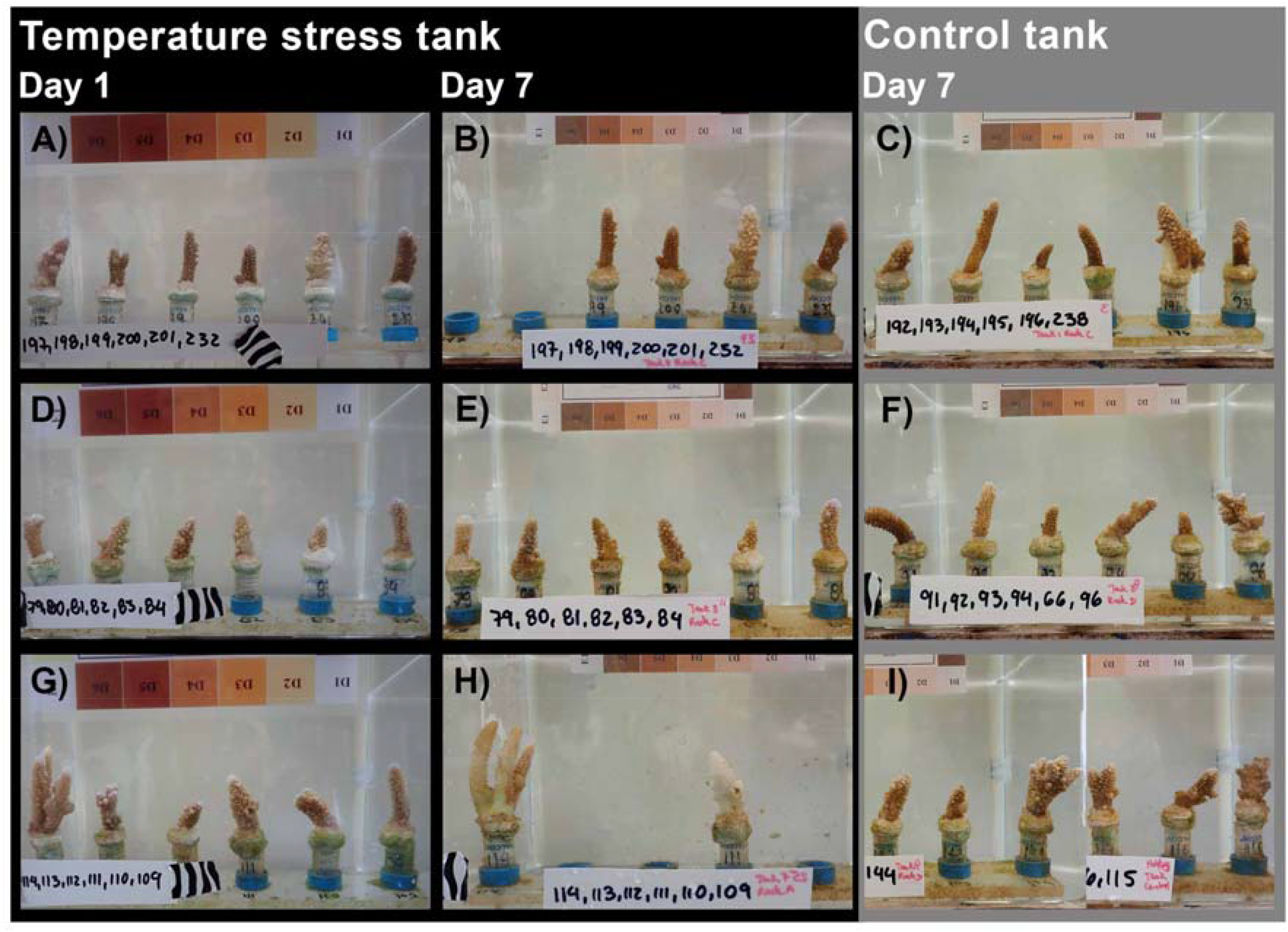
Images of fragments of *Acropora digitifera* taken on day 1 (A, D and G) and on day 7 (B, E and H) of exposure in the temperature stress and the control (C, F, I) treatments. Fragments were removed from the tank on the day they were assessed as dead.

### b) Coral spawning

In anticipation of *A. digitifera* spawning in Palau (Penland et al., 2003; Gouezo et al., 2020), the 34 colonies used in the temperature stress assay were surveyed to assess reproductive (gravid or non-gravid) and health (alive, partial mortality or dead) statuses just before the April full moon (01/04/2018). Reproductive status was established by fracturing two branches per colony and checking for the presence of visible pigmented oocytes (Fig. 3A) (following Baird et al., 2002). Of the eleven colonies identified as RHHT and four colonies identified as RLHT (see above methods and results below), five and three colonies respectively contained visible, pigmented gametes (SM. 2). Three gravid colonies were haphazardly chosen from each RHT category and collected on 29 March for the SB crosses and transported in 50 L containers to PICRC. Colonies were maintained in an outdoor flow-through 760 L holding tank where water was mixed using three magnetic pumps (Pondmaster 1200 GPH). Four days after the full moon, setting (gamete bundles visible within polyp mouths) was observed in all colonies. We used standard coral larval rearing methods (Guest et al., 2010), with modifications to ensure that individual crosses were isolated. From sunset onwards (19:00 h), colonies were checked visually for signs of bundle setting every 30 minutes. As soon as one colony was seen setting, all colonies were isolated in individual 80 L static tanks to prevent cross fertilization. As soon as most bundles were released (Fig. 3B), 200 ml plastic cups were used to scoop buoyant gamete bundles from the water surface. Egg-sperm bundles were separated by transferring them onto a 100 μm mesh filter immersed in a bowl containing a small amount of UV-treated (Trop UV Steriliser Type 6 /IV – TPE, Trop-Electronic GMbH, Germany) 0.2 μm filtered sea water (FSW). Sperm remained in the bowl while eggs remained immersed in FSW within the filter. The filter was removed quickly and transferred to a new bowl with UV-treated 0.2 μm FSW, and eggs were washed five times to remove any sperm residue by carefully adding water into the filter and letting the bowl overflow. Throughout this process, all bowls, filters and other utensils were rinsed with diluted bleach (1%) and FSW. All implements were labelled and used exclusively for individual colonies or crosses to avoid cross contamination. After spawning, colonies were returned to the holding tank, and a week later they were transplanted at the natal reef (Mascherchur).

**Figure 3:**
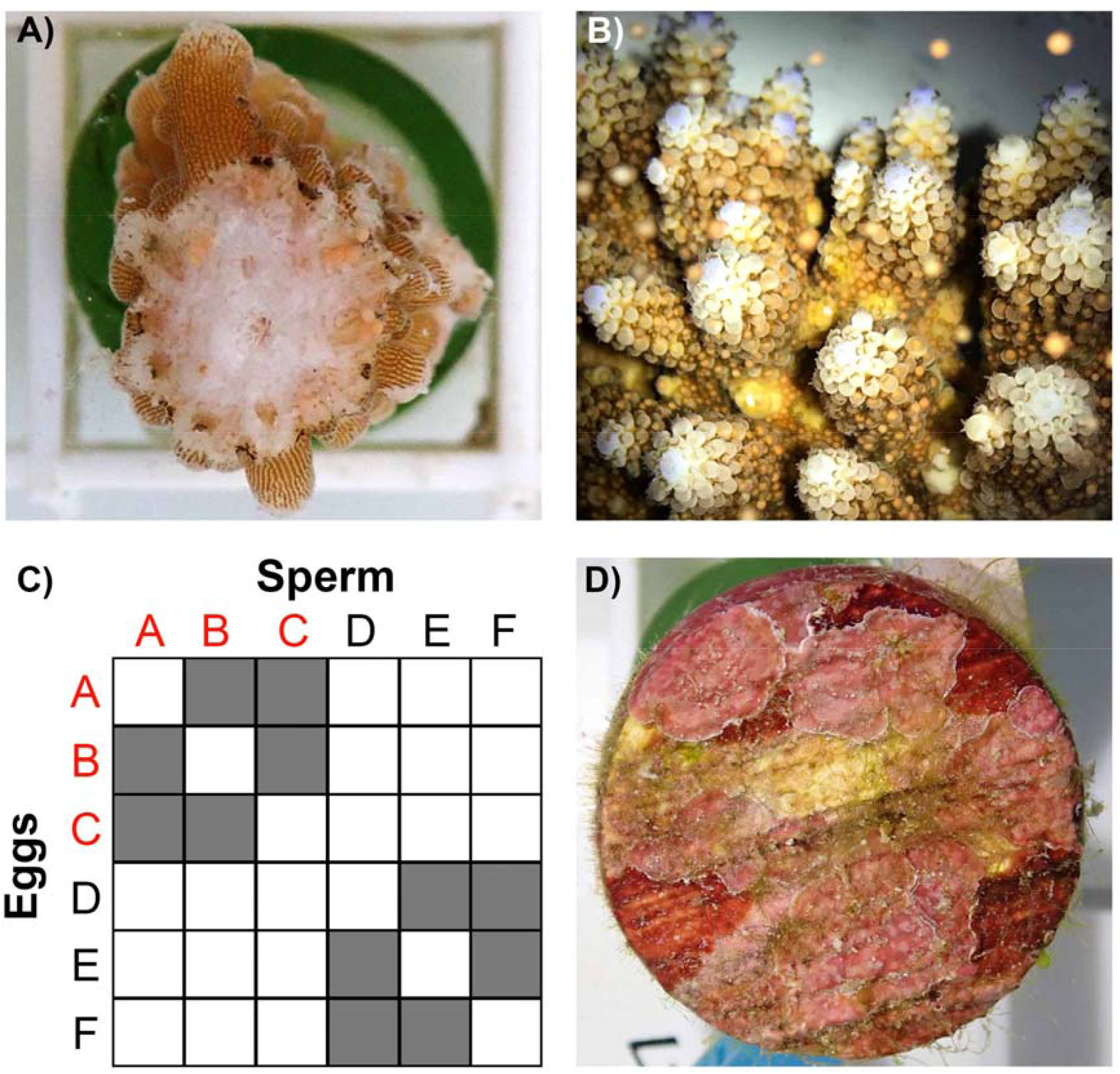
A) Fragment of a gravid colony of *Acropora digitifera* with pigmented eggs. B) *A. digitifera* bundle release. C) Schematic representation of the 12 selective breeding crosses performed using three parental colonies of *A. digitifera* with relative high heat tolerance (RHHT, colonies A, B and C), and three colonies with relative low heat tolerance (RLHT, colonies D, E and F). D) Settlement substrate conditioned with CCA before been offered to larvae for settlement.

### c) Fertilization and selective crosses

Separated gametes were cross fertilized to produce two types of crosses 1) RHHT sire × RHHT dam, and 2) RLHT sire × RLHT dam, and each type of cross was replicated six times using different combinations of parental colonies to produce 12 unique crosses (Fig. 3C). The resulting crosses were maintained in 15 L cone-shaped tanks (Pentair Vaki Scotland Ltd) at ambient temperature, with 0.2 L/min flow-through with UV-treated 0.2 μm FSW, resulting in one turnover per hour per tank (Fig. 4, SM. 3). A PVC “banjo” with a wedge shape, covered with 100 μm mesh filter was fixed to the inside of the outflows of the tanks to avoid loss of larvae. Each cross was divided between two rearing tanks, resulting in 24 larvae culture tanks. Gentle aeration was introduced 24 h after fertilization, when embryo development had progressed sufficiently, and larvae were round and motile.

**Figure 4:**
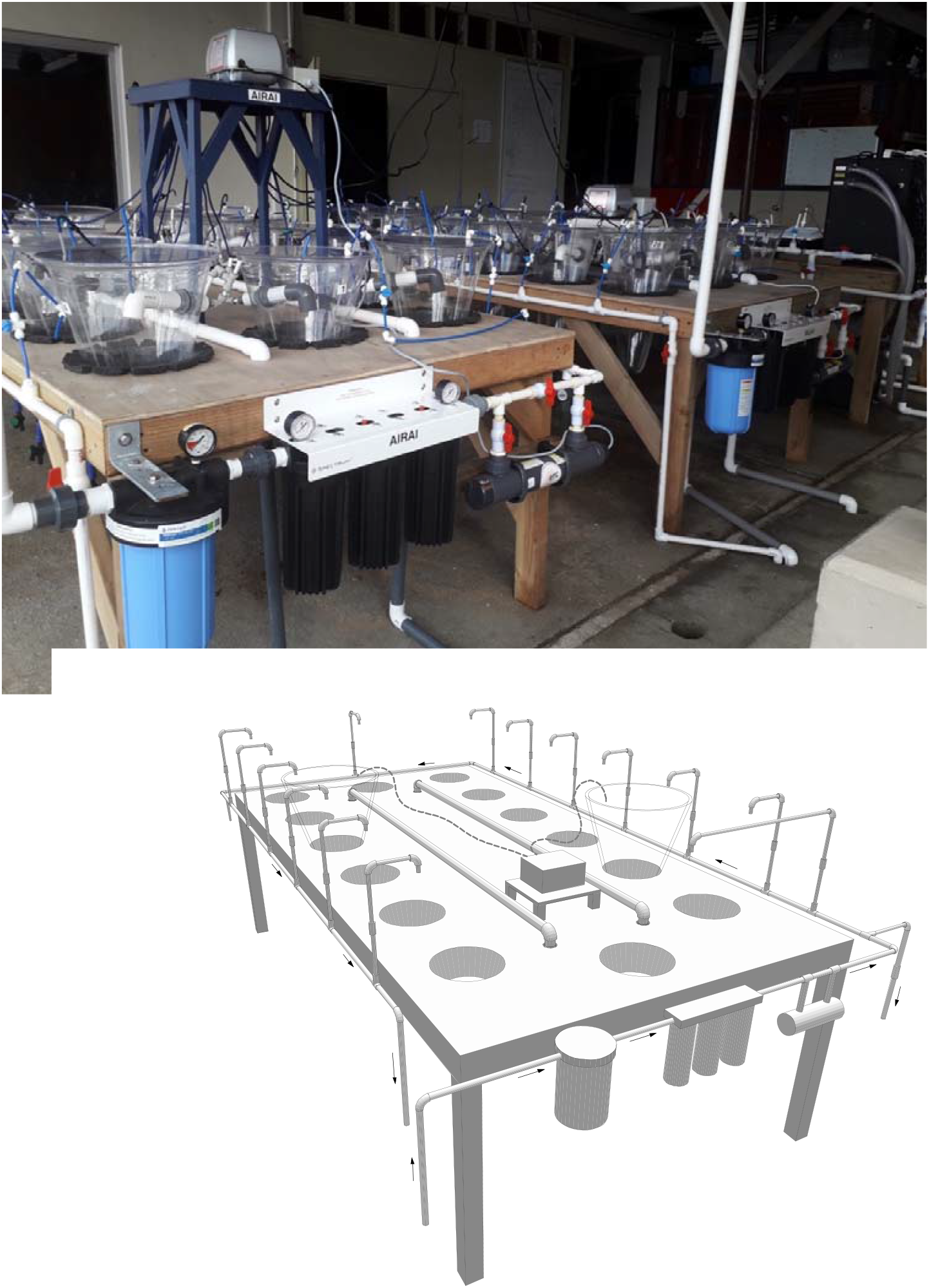
A) Overview of the larval rearing system consisting of 15 L cone shaped tanks with flow-through. Sea water is filtered through 4 filters (50, 10, 5 and 0.2 μm) and then exposed to UV light before entering into the tanks. Each tank has inflow pipe for the filtered and UV treated sea water, an outflow pipe for the waste water, and an airline connected to an air pump. B) Diagram showing the main components of the larval rearing system. Arrows indicated water flow. a) Unfiltered sea water inflow, b) polyvinyl chloride (PVC) valve 1”, c) 1” PVC pipe, d) first filtering station (50 μm), e) 3/4” pipe, f) Second filtering station (10, 5, 1 or 0.2 μm), g) 1/2” pipe, h) Trop UV Steriliser, i) PVC valve 1/2”, j) 15 L cone-shaped tanks, k) Banjo with 100 μm mesh filter, l) Water inflow tube 1/4”, m) John Guest elbow 1/4”, n) John Guest valve 1/4”, o) Water outflow tube 1/2”, p) Air pump, q) Airline, and r) 1” PVC waste pipe.

### d) Larval settlement

Circular ceramic substrates (Oceans Wonders LLC) ∼2 cm in diameter with a 1.5 cm stem (hereafter referred as Substrate Units “SUs”), overgrown with crustose coralline algae (CCA, Fig. 3D) were offered to the larvae for settlement. The SUs had been biologically conditioned for four months (130 days) before spawning in two 300 L holding tanks with flow-through water mixed using four pumps each (Taam Rio +800 Powerhead). SUs were arranged on plastic egg crates raised from the bottom of the tank, with fragments of CCA collected at Mascherchur placed on top of the SUs. To stimulate growth of CCA over the SUs and avoid the colonization of filamentous algae, frames with shading cloth were placed over the tanks to reduce light levels to 4 μmol photons m^−2^ s^−1^. Three days after fertilization, larvae were transferred from 24 larval rearing tanks to 24 settlement tanks filled up with 10 L of 10 μm FSW and 80 conditioned SUs each. Half of the water in each tank was changed daily with new FSW. Two days after larvae were moved to the settlement tanks, SUs with newly settled corals were transferred to flow-through nursery tanks (*ex situ* nurseries). Each SU was tagged with a cable tie using a colour coded system to identify from which cross and replicate culture the recruits had originated (resulting in 24 colour codes from 12 crosses).

### e) Ex situ nursery tanks

SUs with settled corals were randomly distributed among four *ex situ* flow-through nurseries consisting of 184 L tanks (length: 128cm, width: 85cm, water level: 17cm) with 50 μm FSW. Each tank was illuminated with two Aquarium lights (48” 50/50 XHO Led, Reef Brite Ltd) at an intensity of 400 μmol photons m^−2^ s^−1^ over a 12 h:12 h diurnal cycle, and had two pumps (Hydor Koralia Nano Circulation Pump/Powerhead, Fig. 5A, B) to create water circulation. Fragments from each parental colony were added to each tank to promote Symbiodiniaceae uptake by the coral settlers, together with fragments of CCA. To minimise growth of turf algae, eight small herbivorous juvenile rabbitfish (*Siganus lineatus*, ∼5 cm long) and numerous small grazing snails (*Cerithium sp*.) were added to each tank. Fish were fed daily with fish pellets (Ocean Nutrition Formula) and nurseries were cleaned every other week by syphoning off the detritus from the bottom of the tanks.

**Figure 5:**
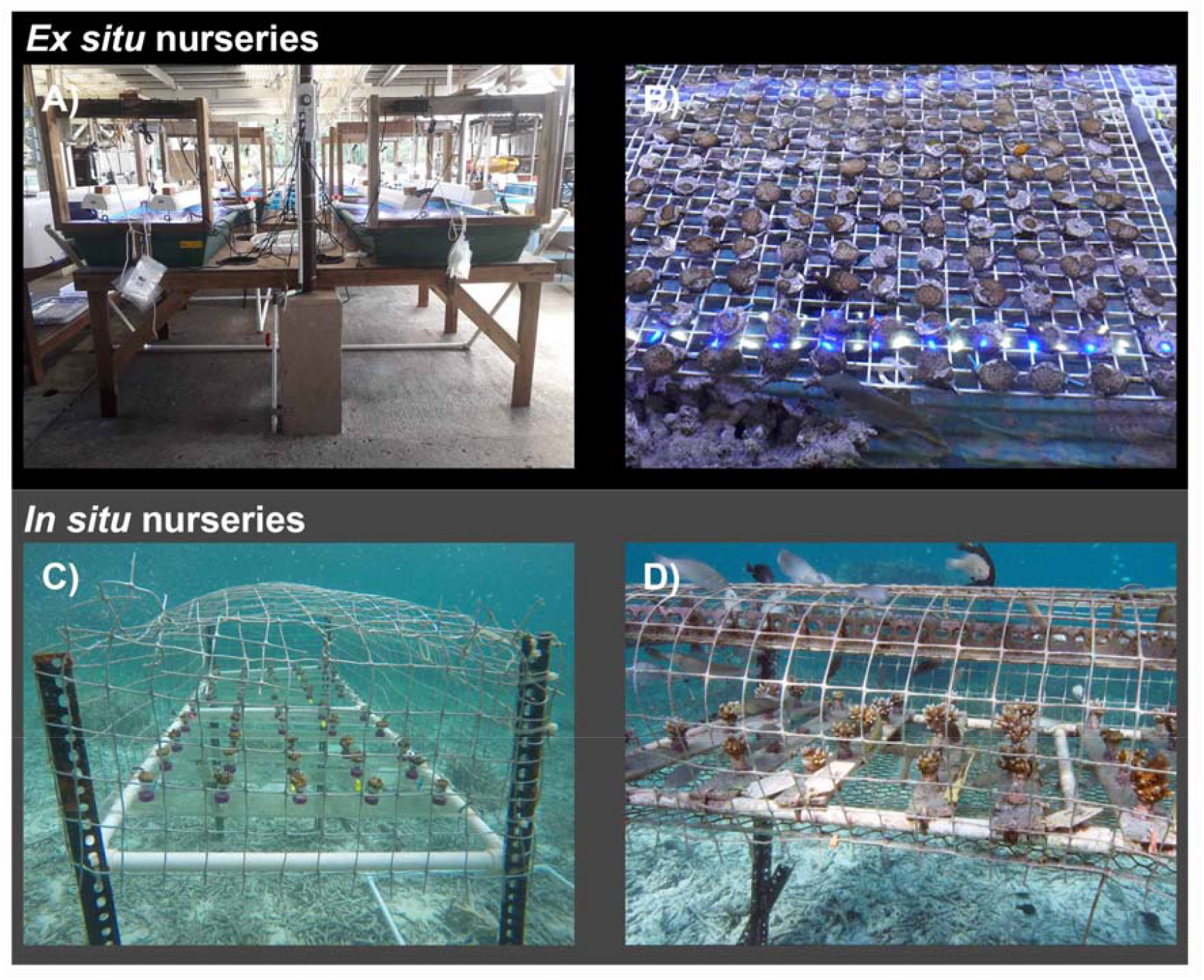
A) *Ex situ* nursery tanks, B) Detail of *ex situ* nurseries with the presence of herbivores *Siganus lineatus*. C) Coral colonies when transferred from the *ex situ* to the *in situ* nursery (13-months old). D) Corals (17-months old) in the *in situ* nursery showing the presence of herbivore fish.

### f) Coral outplanting to the reef

Coral colonies from 12 crosses were outplanted from the *ex situ* nursery to the natal reef of parental colonies (Mascherchur) when colonies were five and 11-months old (144 and 318 days, n = 288 and 96 colonies respectively, Table 1). The number of colonies outplanted at 11 months was limited by the workforce available at the time. To facilitate monitoring, SUs with corals were outplanted along 16 fixed 10-m transects at a depth interval of 1.5 to 4 m. To attach each SU to the reef substrate, 11 mm holes were drilled into bare reef substratum with a submersible cordless drill (Nemo Divers Drill) and SUs were glued with epoxy (Milliput Standard) after cleaning the reef surface area with a wire brush. Nails were hammered next to each SU to which a cable tie, colour coded for each cross was attached. Two divers were required to outplant 25 corals in one 2-hour dive. Colonies were monitored at 11, 17, 25, and 32 months (318, 515, 767, and 974-days old respectively) to assess their status (alive, missing or dead) and photographed from directly above with an underwater camera (Olympus Tough TG-5), and with a ruler for scale.

**Table 1:** Number of colonies from the selective breeding crosses outplanted to Mascherchur at two different ages.

| Parental colonies<br>relative heat<br>tolerance | Parental<br>colonies | Five-months old | 11-months old |
| --- | --- | --- | --- |
| RHHT | A x B | 30 | 8 |
|  | A x C | 26 | 10 |
|  | B x A | 17 | 7 |
|  | B x C | 27 | 8 |
|  | C x A | 11 | 2 |
|  | C x B | 8 | 4 |
|  | C x B | 12 | 6 |
| RLHT | D x E | 25 | 8 |
|  | D x F | 28 | 9 |
|  | E x D | 32 | 8 |
|  | E x F | 27 | 6 |
|  | F x D | 19 | 8 |
|  | F x E | 26 | 8 |
| Total |  | 288 | 92 |

### g) In situ nurseries

After 13 months (386-days) of *ex situ* rearing, the remaining colonies on 296 SUs were transferred to six *in situ* nurseries (N 7°18’19.80”N; E 134°30’6.70”E, Fig. 1B) 2.20 km away from the natal reef. Nurseries were constructed *in situ* using steel slotted angle bars 40 × 40 mm (length: 135 cm, width: 60 cm), raised from the seafloor (85 cm). A plastic mesh (aperture size 5 cm) was used to cover the nursery structures (Fig. 5C, D) to exclude larger corallivores (Baria et al., 2010). Corals were attached to plexiglass racks with five colonies per rack, spaced 3 cm apart. Ten racks were placed in each nursery resulting in a total of 50 colonies per nursery. Meshes and racks in each nursery were cleaned monthly using a stiff plastic brush to remove algal overgrowth. When colonies were 17-months old (504 days), each colony was photographed with an underwater camera (Olympus Tough TG-5) from directly above, and with a ruler for scale.

### h) Costs of producing, rearing, outplanting and monitoring ∼2.5 years old coral colonies using a selective breeding approach

The cost of producing ∼2.5-years old (32-months) live colonies was calculated from the total cost of materials and hours of labour needed to run the experimental setup at full capacity with 24 larval cultures, rear, outplant at the natal reef and monitor the resulting colonies. The cost of the experiment to characterize the RHT of the parental colonies was not included in this analysis, as this will vary considerably according to the trait of interest being selected for, the methodology (i.e., experiments under laboratory conditions, SNP analysis, etc.) used to identify colonies of interest, and their location. Due to logistical constraints, we were not able to quantify fertilization success, larval survivorship and initial settlement densities and survivorships, hence values reported in the literature were used for the analysis. Details of the assumptions of the analysis, costs of consumables, equipment and person hours are provided in the Supplementary Material (SM. 4). Cost per coral was estimated by dividing the total cost for the project by the number of SUs containing one surviving 2.5-year old coral for different ages at outplant (five or 11 months). To compare costs of rearing in *ex situ* and *in situ* nurseries we considered the costs of consumables for their construction and their maintenance during a 11-month period, and cost per SUs was estimated by dividing the total cost of building and maintenance of the nurseries by the number of SUs. The overall efficiency for each outplantation age (five or 11 months) was estimated by dividing the number of coral eggs used by the number of living colonies after 2.5-years. The aim of this analysis was to: 1) estimate the minimum cost of CLP using a SB framework, 2) identify the steps of the framework (coral collection, spawning to competency, settlement, rearing, outplanting and monitoring) that incur the highest cost of the budget, and 3) evaluate the effect on efficiency and cost per coral of outplanting colonies at two ages (five and 11 months). The total cost of this framework should not be used as a reference for SB under AE since: 1) specific assumptions were made for its computation (SM. 4) and any change in these assumptions will change the total costs, 2) upscaling each of the steps will reduce their cost due to economies of scale, 3) our results do not include data on the reproductive status of the recruited colonies 2.5 years post-fertilization. Before the predicted spawning event in 2020, the reproductive status of 9 colonies with the biggest diameter in the *in situ* nursery was assessed (as described in section b), with none containing visible eggs (SM. 6). Additionally, recruited colonies had smaller sizes than the corals in the *in situ* nursery and were only starting to develop branches (Fig. 6), suggesting that most probably they were not reproductive at the time of spawning, however we have no data to support this.

**Figure 6:**
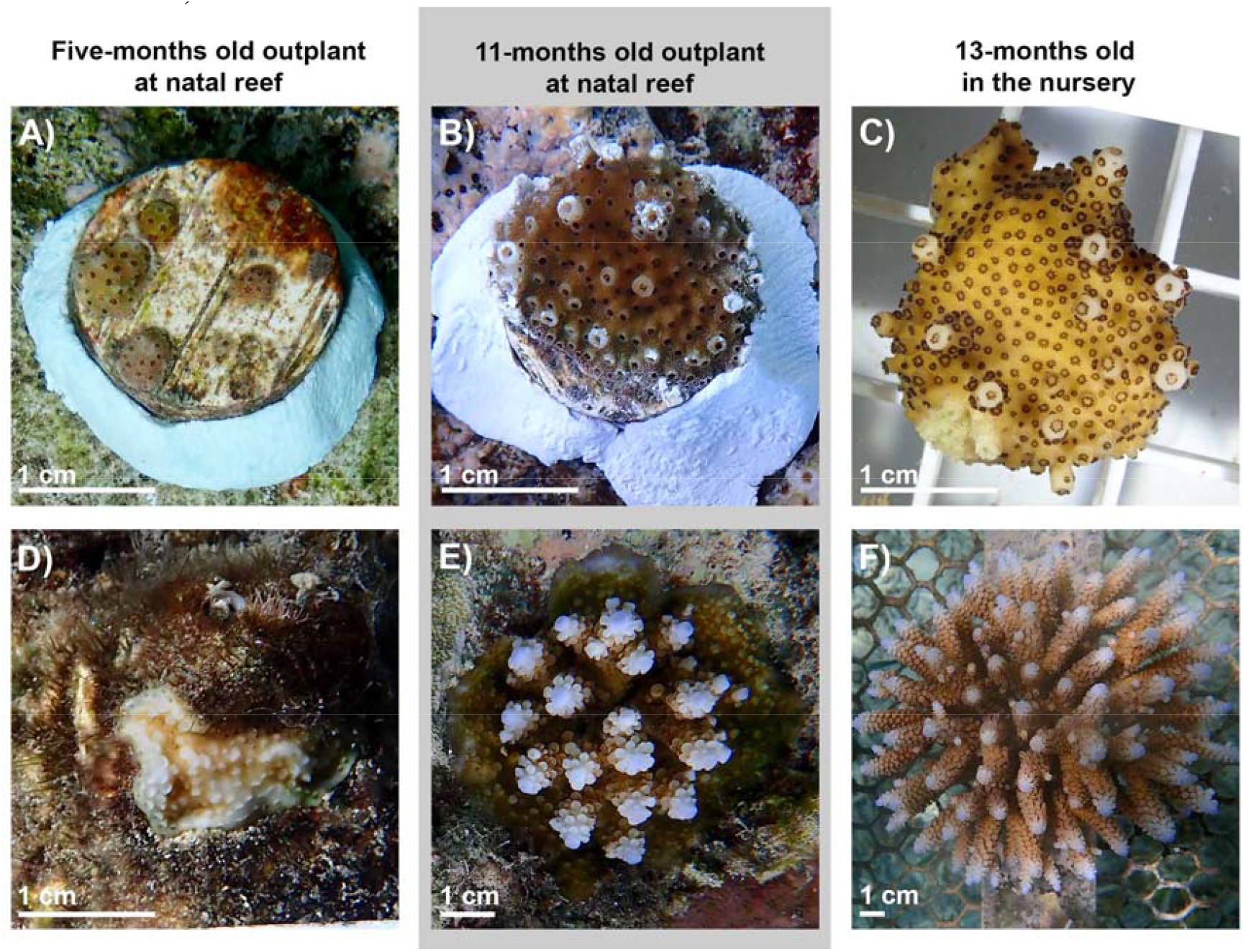
Images showing representative colonies to illustrate differences in size and morphology at the time of outplanting to the reef, or transfer to the *in situ* nursery (A: five-month old outplant, B: 11-month old outplant and C: 13-month old before transferring to the *in situ* nursery), and during the last monitoring (D: five-month old outplant, E: 11-month old outplant and F: *in situ* nursery) when colonies were 2.5 years old (52 days after the spawning event of 2020).

### Settlement density

To provide practical guidance for planning sexual coral propagation programmes, the minimum number of settlers needed to obtain SUs with at least one colony after four months (132-days old) of nursery rearing was determined by testing the effect of settlement density on colony survivorship. This experiment was carried out during the spawning event of the 6^th^ of April 2020 using three adult *A. digitifera* colonies collected at Mascherchur. A mass culture was produced with the gametes of the three colonies following the protocol previously described for collecting bundles and rearing larvae. Four days after spawning, once larvae were competent to settle, 1.5 L static tanks were stocked at three levels of larval densities (10, 25 and 50 larvae per SU or 67, 167 and 333 larvae L^-1^, with n = 8 replicate tanks). Each tank contained ten SUs previously conditioned with CCA for 198 days (SM. 7). Water changes were carried out twice daily over a week with UV treated 1 μm FSW. Ten days after settlement, the number of settlers on each SUs were counted using a stereomicroscope. SUs (n = 157) with live settlers (between one and 15 per SU) were then randomly distributed across four *ex situ* nurseries (described in section e). The number of live corals on each SU was again recorded after four months using a stereomicroscope.

### Data analysis

Natural mortality of tagged adult colonies on the reef was estimated using yearly exponential rates of survival (Clark and Edwards, 1995). Colony survivorship was compared between the two different outplanting times to the reef (five and 11-month old) using right censored data with the Kaplan–Meier model with the log-rank statistic (Harrington and Fleming, 1982). As it was not possible to determine the exact time of death for each coral, the date that a coral died was estimated as the middle time point between survey dates. Survival functions of corals were compared (a) among outplanting times at different ages (five and 11-months old), and (b) once outplanted i.e. with respect to days out on the reef rather than age attained. Colony size, measured as planar area for corals outplanted to the reef (at an age of five and 11-months) or moved to the *in situ* nursery was estimated from scaled downward-facing images taken when corals were 17-months old using Image J. During image analysis, the number of distinctive developed branches per colony was recorded as an indicator of volume. The number of colonies with branches was compared using basic descriptive summary methods (percentages of branching colonies of the total alive outplants). The effect of outplanting method (3-level fixed effect) on colony size at 17 months was tested using Generalized Linear Mixed effects Models (GLMM, Brooks et al., 2017), accounting for differences in size due to cross replication (12-level random factor). Log-Gamma link was used to prevent negative fitted values of this strictly positive response variable. Among method comparisons were tested using a post-hoc Tukey Test.

The effect of larval culture density (3-level fixed effect) on settlement density after 10 days was tested using GLMM, accounting for variability among settlement tanks (8-level random effect). The relationship between settlement density (fixed effect) and number of four month-old colonies per SU was tested using GLMM, accounting for variability among holding tanks (4-level random effect). All model validation steps included assessing homogeneity of residuals versus fitted values, over and under dispersion and a simulation study to test the ability of the model to capture zero-inflation (Zuur and Ieno, 2017). Poisson models suitable for count data (Bates et al., 2015) were over-dispersed due to underestimation of zeros, however, negative binomial models with quadratic parameterization variance structure (Brooks et al., 2017) passed all validation routines (SM. 8). Marginal R^2^ values, the proportion of variance explained by fixed effects alone were computed for the final model (Lüdecke et al., 2020).

## Results

### Selection of parental colonies

RHT varied among colonies (n = 34), with 32% categorised as RHHT and 15% categorised as RLHT (Table 2). For 21% of the colonies at least one of the fragments held in the ambient control tanks died, and therefore their RHT was considered as unresolved (Table 2) and for the remaining 32% colonies, between 25% and 66% of fragments died during the experiment and colonies were unclassified in terms of RHT (Table 2). Natural mortality rates of adult tagged colonies at Mascherchur were estimated at ∼20% per year, with eight colonies recorded as dead 140 days after tagging (n = 99). Partial mortality was observed in 6% of the tagged colonies and one colony could not be relocated (SM. 2). From the surviving, visibly healthy colonies (n = 84), 38% contained pigmented eggs in March 2018, and these included five colonies classified as RHHT and three as RLHT (SM. 2).

**Table 2:**
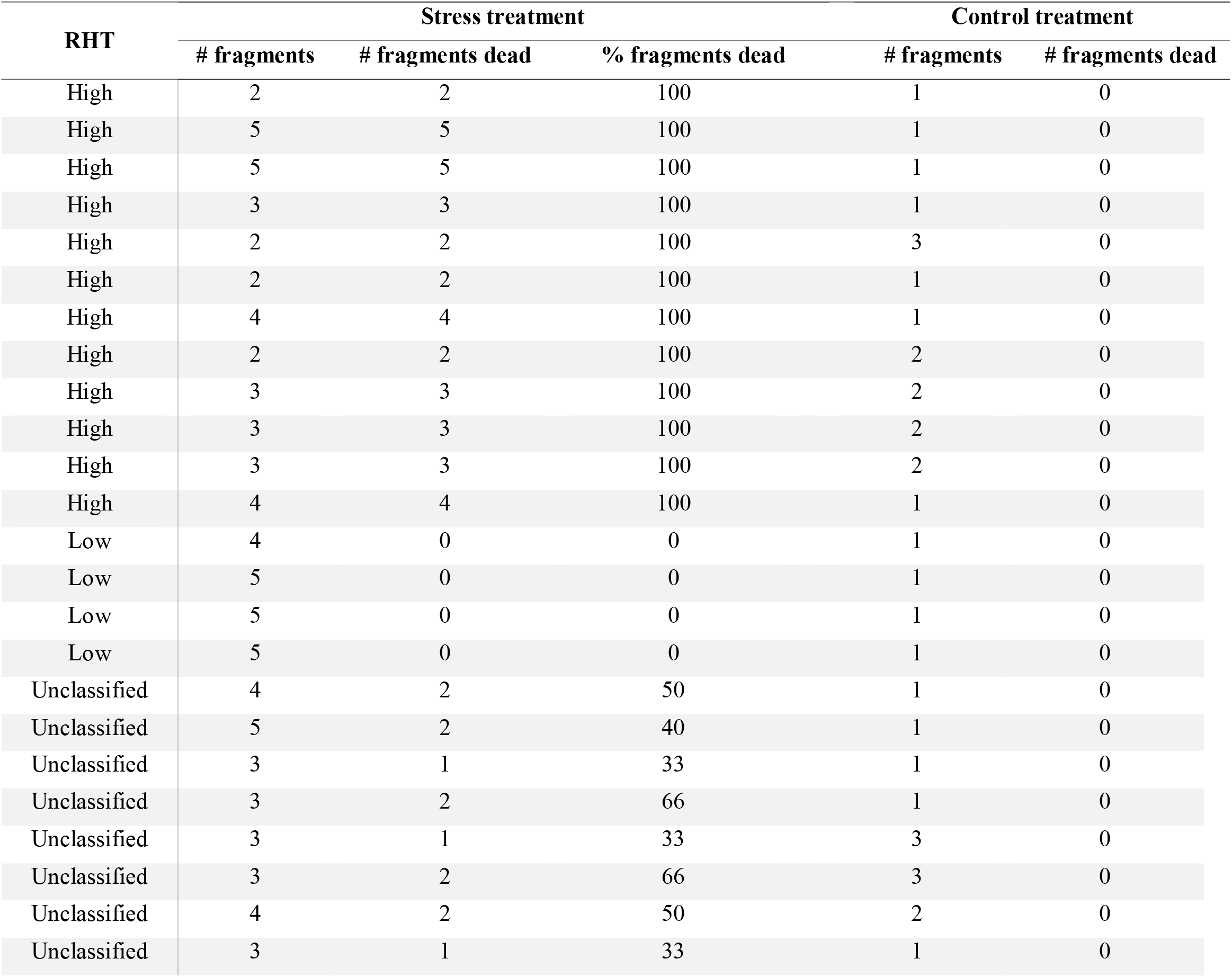
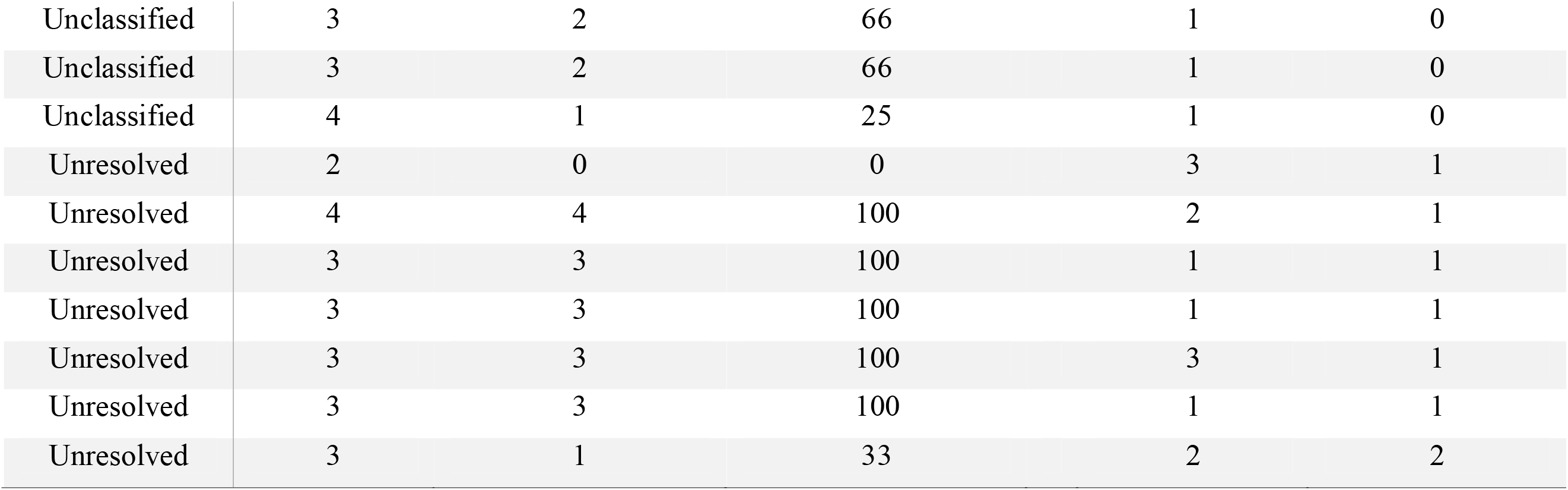
Results of the 7-day heat stress exposure used to determine the relative heat tolerance (RHT) of the colonies.

| RHT | Stress treatment |  |  | Control treatment |  |
| --- | --- | --- | --- | --- | --- |
|  | # fragments | # fragments dead | % fragments dead | # fragments | # fragments dead |
| High | 2 | 2 | 100 | 1 | 0 |
| High | 5 | 5 | 100 | 1 | 0 |
| High | 5 | 5 | 100 | 1 | 0 |
| High | 3 | 3 | 100 | 1 | 0 |
| High | 2 | 2 | 100 | 3 | 0 |
| High | 2 | 2 | 100 | 1 | 0 |
| High | 4 | 4 | 100 | 1 | 0 |
| High | 2 | 2 | 100 | 2 | 0 |
| High | 3 | 3 | 100 | 2 | 0 |
| High | 3 | 3 | 100 | 2 | 0 |
| High | 3 | 3 | 100 | 2 | 0 |
| High | 4 | 4 | 100 | 1 | 0 |
| Low | 4 | 0 | 0 | 1 | 0 |
| Low | 5 | 0 | 0 | 1 | 0 |
| Low | 5 | 0 | 0 | 1 | 0 |
| Low | 5 | 0 | 0 | 1 | 0 |
| Unclassified | 4 | 2 | 50 | 1 | 0 |
| Unclassified | 5 | 2 | 40 | 1 | 0 |
| Unclassified | 3 | 1 | 33 | 1 | 0 |
| Unclassified | 3 | 2 | 66 | 1 | 0 |
| Unclassified | 3 | 1 | 33 | 3 | 0 |
| Unclassified | 3 | 2 | 66 | 3 | 0 |
| Unclassified | 4 | 2 | 50 | 2 | 0 |
| Unclassified | 3 | 1 | 33 | 1 | 0 |
| Unclassified | 3 | 2 | 66 | 1 | 0 |
| Unclassified | 3 | 2 | 66 | 1 | 0 |
| Unclassified | 4 | 1 | 25 | 1 | 0 |
| Unresolved | 2 | 0 | 0 | 3 | 1 |
| Unresolved | 4 | 4 | 100 | 2 | 1 |
| Unresolved | 3 | 3 | 100 | 1 | 1 |
| Unresolved | 3 | 3 | 100 | 1 | 1 |
| Unresolved | 3 | 3 | 100 | 3 | 1 |
| Unresolved | 3 | 3 | 100 | 1 | 1 |
| Unresolved | 3 | 1 | 33 | 2 | 2 |

### Selective breeding

A total of 12 crosses, each replicated twice, were performed using gametes from six donor colonies. The collection and separation of gametes, the performance of SB crosses, the washing of embryos after fertilization, and the maintenance of cultures were carried out by six researchers, two of them with expertise in *ex situ* coral spawning (JG and AH). Embryos were reared in all replicate cultures and larvae were successfully settled onto 1920 SUs.

### Settlement density experiment

In 2020, the density of the larval culture had a significant effect on settlement densities (Fig. 7A). The expected mean settlement density using a larval culture of 50 larvae per SU was two- and three-fold higher than that of the two other larval cultures with 25 and 10 larvae per SU respectively (GLMM, R^2^ = 0.28, P < 0.01, SM. 9). The density of live settlers per SU at 10 days post-settlement had a positive effect on the density of colonies four months later (Fig. 7A, GLMM, *R*^*2*^ = 0.36, *P* < 0.001, SM. 10). On average, a settlement density of four settlers per SU was sufficient to obtain at least one coral per SU after four months under *ex situ* nursery conditions (1.3 settlers per cm^2^ of effective settling surface, Fig. 7B).

**Figure 7:**
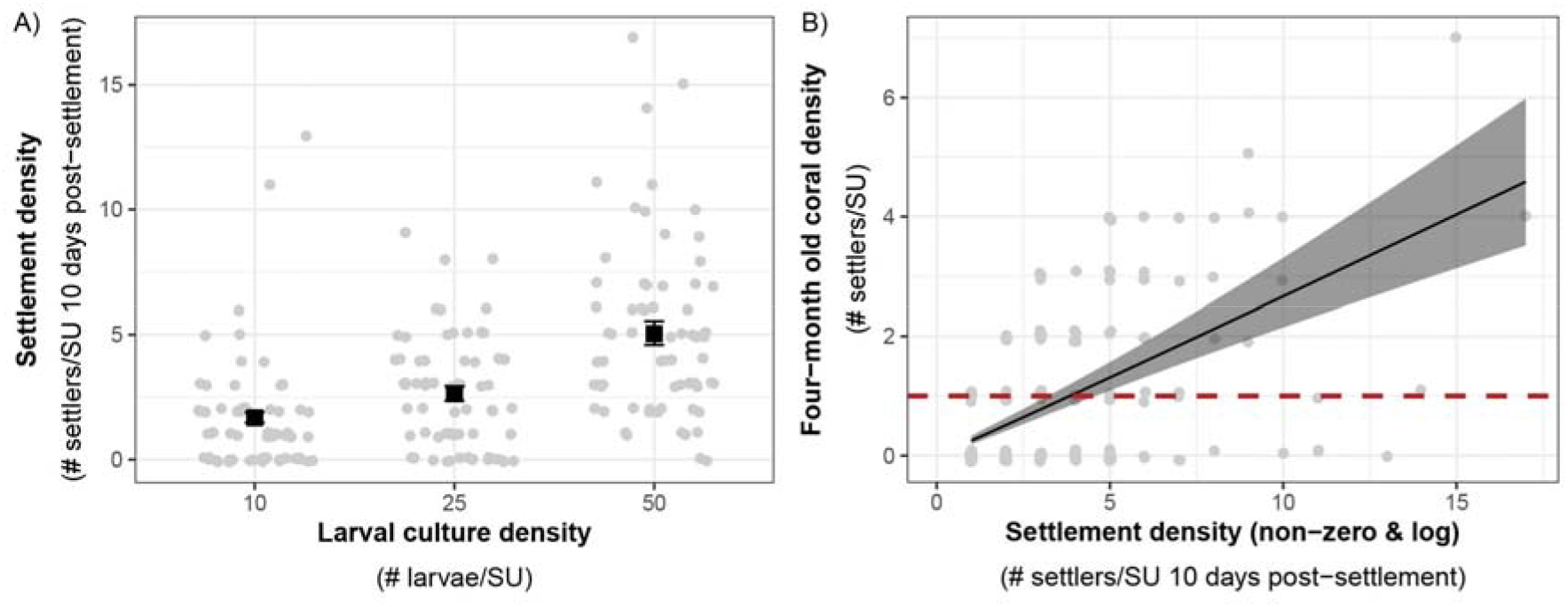
A) Effects of larval density on settlement density. Average expected response values among settlement tanks (black squares) from negative binomial GLMMs are shown with standard error bars. B) Expected relationship between settlement density and settler density after four months (132 days, black line and standard error) among *ex situ* nursery tanks derived from a negative binomial GLMM. Equation: *four months coral density = exp(−1*.*44 + 1*.*02* * *log(Non-zero settlement density))*. The critical target threshold of 1 coral per substrate after four months is represented with a red dashed line. Raw count data (grey circles) are visualised with a jitter to minimise overlap of points in the 2 graphs.

### Effects of ex situ and in situ nursery rearing on colony survivorship

Colony survivorship was significantly affected by age at the time of outplanting (F_1,380_=102, p < 0.05, Fig. 8). The median survival age of corals outplanted to the reef at 11-months old was more than twice that of those outplanted at five-months old (median survival age of 646 and 257 days respectively). The relative survival once outplanted (i.e. with respect to days out on the reef) increased more than threefold when *ex situ* nursery time was extended, with colonies outplanted five months post-fertilization surviving a median time of 104 days, whereas outplants at 11-months post-fertilization survived for a median time of 324 days post-outplant. Only ∼6% of corals that were outplanted at five months post-fertilisation survived to 32-months old. In contrast, corals that were reared in nurseries for 11-months prior to outplanting to the reef had five times better survivorship (∼30%) to 32-months old.

**Figure 8:**
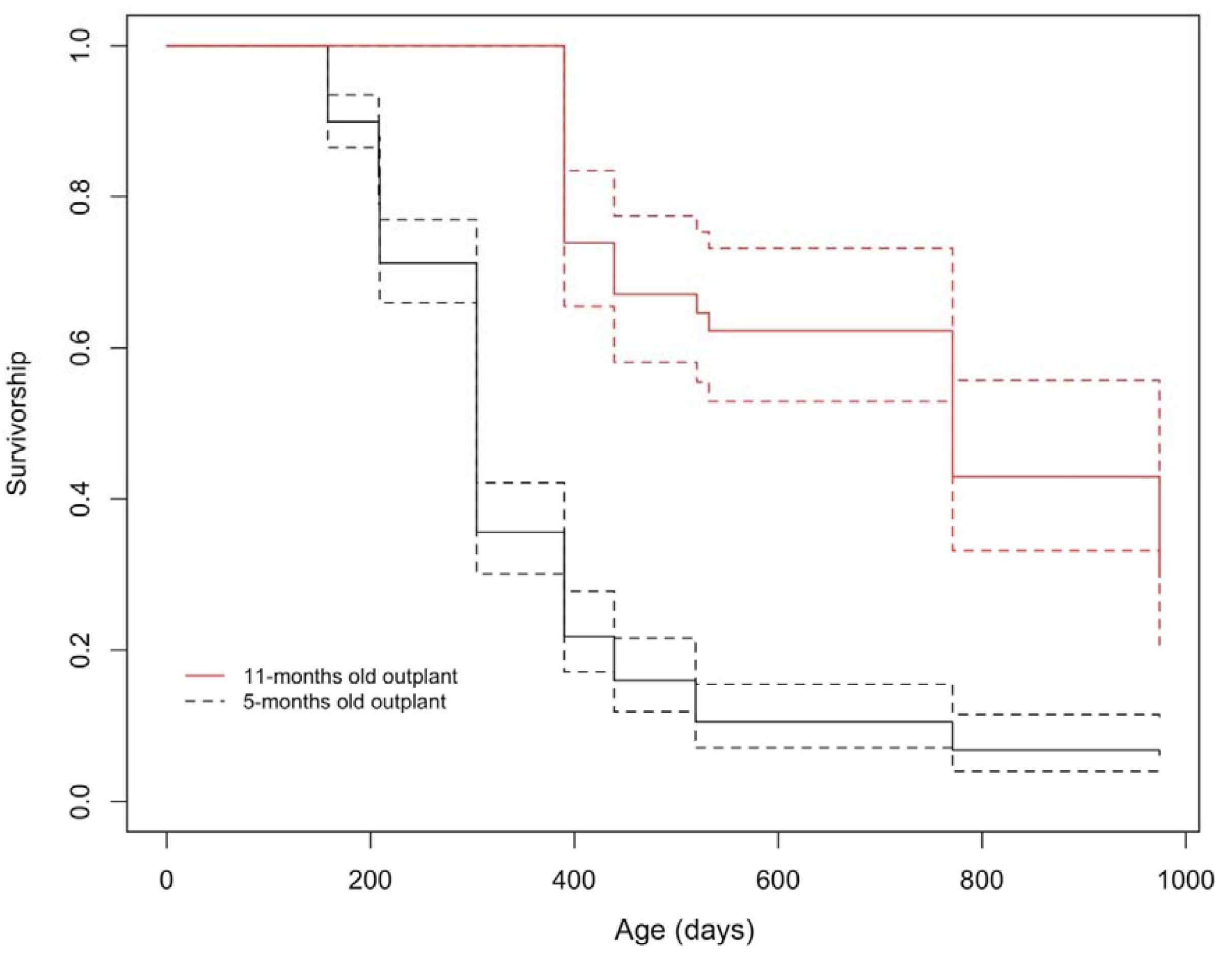
Survivorship curves for *Acropora digitifera* corals outplanted to the reef when colonies were 5-months (black line) and 11-months old (red line).

At 17 months, corals at the *in situ* nursery started developing a 3D structure with branches present in 72% of the colonies, compared to corals outplanted to the reef where none had started branching for both five and 11-months outplants. The planar area of corals was greater in treatments that had longer husbandry times (i.e. were held in the *in situ* or *ex situ* nurseries for longer periods). Coral size at 17 months was strongly affected by age at time of outplanting (i.e. outplanted to the reef at five month, 11 months, or transferred to the *in situ* nursery after 13 months Fig. 9). *In situ* nursery reared corals were significantly larger than those outplanted at five and 11-months (GLMM p < 0.001, SM. 11), and corals outplanted at 11-months were significantly larger than those outplanted at five months (GLMM Tukey Test, p < 0.001).

**Figure 9:**
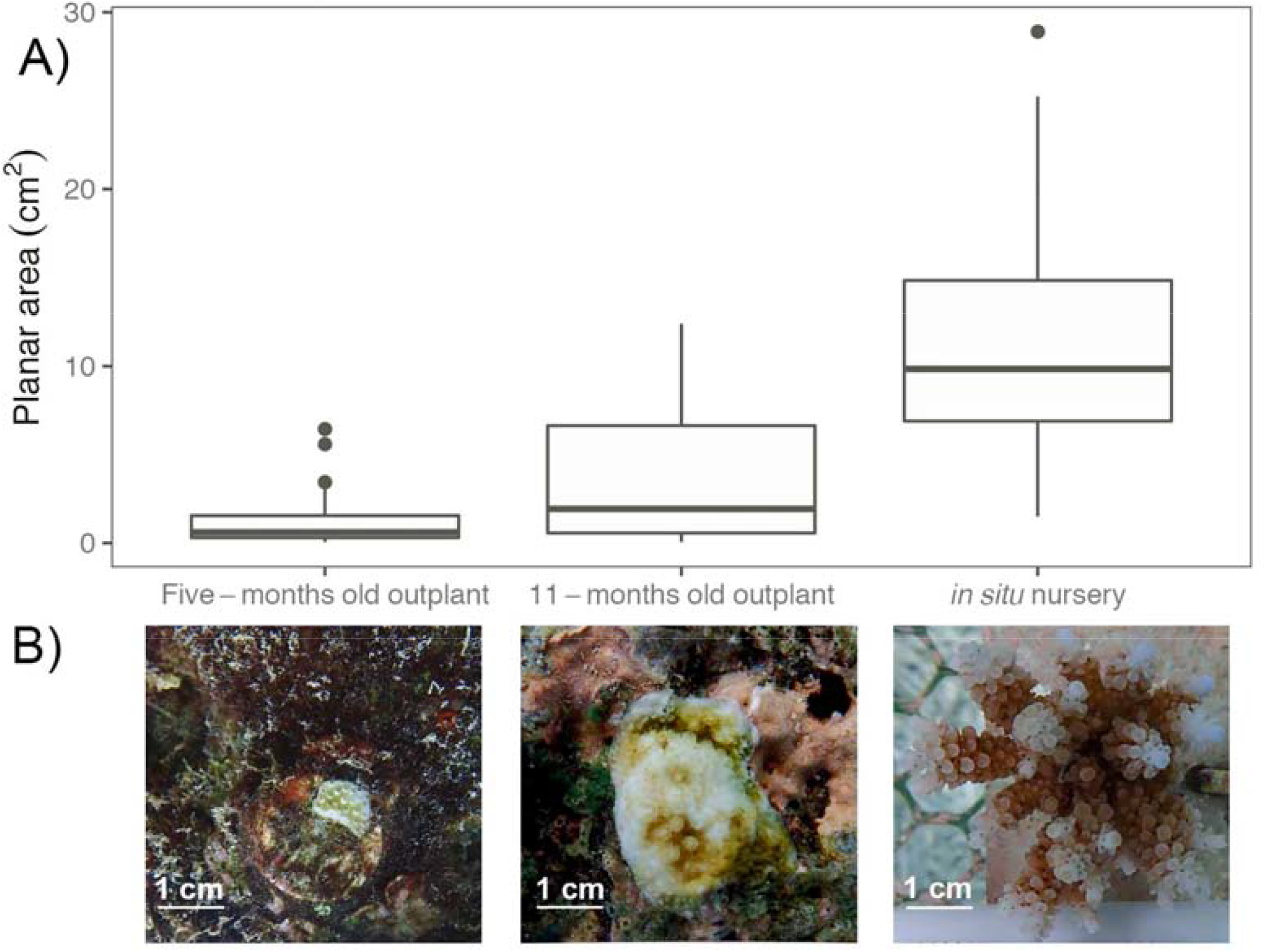
A) Planar area comparison between colonies from the *ex situ* nursery that were outplanted to the reef at five and 11-months old or transferred to the *in situ* nursery at 13-months old. B) Images showing representative corals in the different treatments at the same age (17-months old).

### Cost analysis

The total cost of producing, rearing, outplanting and monitoring 2.5 years old corals using a SB framework was US$23,817 for colonies outplanted at five-months old (SM. 12, SM. 13), and US$22,500 for those outplanted at 11-months old (Table 3, SM. 14). The overall efficiency to produce 12 distinctive crosses with two replicate cultures each and rearing the resulting colonies under *ex situ* nursery conditions until outplant age (either five or 11-months old) was increased fourfold when outplanted at the later age (0.09 and 0.4 respectively). Equally, the cost of each SU with a live coral after 2.5 years was five times lower when nursery period was extended (US$227 for outplants at five-months old compared to US$49 for 11-months old outplants). Outplanting was the activity that incurred the highest costs (35% and 32% of the total cost for the five and 11-months old respectively; Table 3, SM. 14), followed by the monitoring (33% and 31% of the total cost for the five and 11-months old respectively, Table 3 SM. 14). Costs associated with rearing corals for 11 months in *ex situ* compared to *in situ* nurseries differed considerably, with cost per SUs being sixfold lower for the *ex situ* nurseries (US$0.97 and US$5.75 respectively, SM. 14, 15, and 16).

**Table 3:** Cost of sexually propagating *Acropora digitifera* corals using a selective breeding framework with corals reared under *ex situ* nursery tanks until 11-months old when they were outplanted to the reef. The capital equipment is pro-rated over five years.

| Category | Capital equipment | Consumables | Labour | Total | % total cost |
| --- | --- | --- | --- | --- | --- |
| Coral collection | \$426 | \$785 | \$312 | \$1,521 | 7 |
| Spawning to competency | \$1,353 | \$803 | \$1,274 | \$3,430 | 15 |
| Settlement | \$181 | \$624 | \$213 | \$1,018 | 5 |
| *Ex situ* nursery rearing | \$1,592 | \$254 | \$575 | \$2,421 | 11 |
| Outplanting at 11-months old | \$347 | \$4,807 | \$2,010 | \$7,164 | 32 |
| Monitoring and maintenance | \$559 | \$3,525 | \$2,861 | \$6,945 | 31 |
| **Grand Total** | **\$4,459** | **\$10,798** | **\$7,243** | **\$22,500** | **100** |

## Discussion

Research to assess the feasibility of assisting adaptation of corals in the face of ocean warming is accelerating, however frameworks for practical application of the outputs of this research to conservation and management are lacking. Selective breeding (SB), one of several proposed assisted evolution (AE) techniques (van Oppen et al., 2015), requires selection of parental colonies prior to their sexual propagation to produce specific crosses. Considerable advances in coral larval propagation (CLP) as a tool for reef rehabilitation have been achieved in recent years, but so far these have used randomly selected colonies with no attempt to select for specific traits in parental colonies. Here we present a framework for CLP that involves prior selection of parental colonies to perform multiple paired crosses based on intrapopulation variation in their heat tolerance (Fig. 10). We established optimal settlement densities of larvae to obtain one coral per SU after four months of *ex situ* rearing and demonstrate the potential of rearing corals using a combination of *ex situ* and *in situ* nurseries to optimise their growth and survivorship. We found large differences in growth between corals outplanted to the reef and those reared in *in situ* nurseries. We also found differences in survivorship depending on age at outplant from *ex situ* nurseries, emphasizing the benefits of long nursery phases on final costs per coral. Our data highlight some of the major challenges associated with combining SB with CLP, and the areas that need further research and development to improve efficiency and reduce the high costs involved.

**Figure 10:**
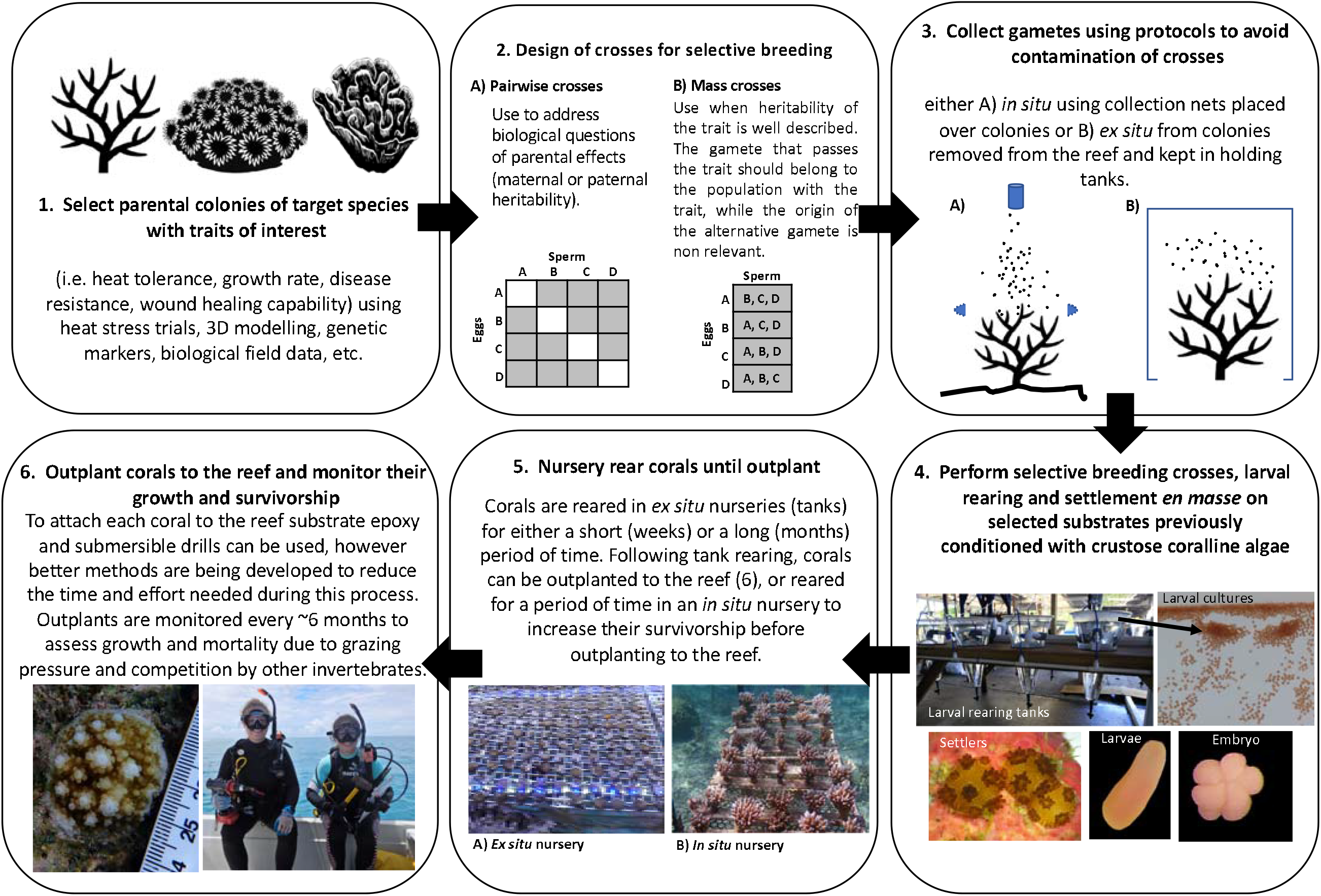
Practical framework for coral reef restoration ecology using selective breeding.

In this study, the selection of parental colonies was made based on intrapopulation variation in heat tolerance using a 7-day temperature stress exposure (Fig. 10.1). Our results show that our study population of *A. digitifera* contained considerable intrapopulation variability in RHT, with relatively high proportions of colonies (∼32%) able to withstand a relatively severe 7-day heat stress with zero fragment mortality. In this case, there was clearly sufficient intrapopulation variability in RHT to provide broodstock for SB initiatives, although further work is needed to determine if the selected trait is passed on to offspring. The high annual mortality rate of adult colonies (9.5%) and the low proportion of the population spawning in the same month (38%) meant that we had relatively few colonies to choose from at the time of doing the selective crosses. Rates of annual mortality recorded here are typical for *Acropora* (Madin et al., 2014) and spawning synchrony can vary considerably among taxa, location and year (Baird et al., 2009; Gouezo et al., 2020), or can shift with handling and transportation stress or while in captivity (Okubo et al., 2006; Craggs et al., 2017). These facts highlight the need to identify and test relatively large numbers of individual corals within populations to increase the likelihood of having sufficient numbers of spawning adults for selective breeding. Selecting the broodstock from within populations (rather than selecting and crossing corals from distinct populations) avoids risks associated with maladaptation, genetic swamping or outbreeding depression (Edmands, 2006; Baums, 2008). It also reduces logistical challenges associated with the collection and transport of colonies or gametes between populations (i.e. between geographically distant locations or countries). Conversely, a limitation of conducting SB within populations is the need to identify sufficient numbers of parental colonies with the desired trait *a priori* using appropriate stress trials. Sufficient numbers of these tagged colonies also need to survive, remain healthy, and be gravid at the predicted spawning time. An alternative to direct testing for a desired phenotype is the identification of biomarkers of stress tolerance (*i*.*e*., a specific lipid or protein constituents, immune profiles, genes, microbial or Symbiodiniaceae symbiont communities, etc.). An option to overcome this hurdle is to create clonal broodstocks of colonies with desired traits or biomarkers either in an *in situ* nursery or at the parental reef. A combination of these two practices could increase the access to genetic material for SB, and even provide access to the same broodstock during consecutive years.

In the present study, we limited our selective breeding design to single pairwise crosses (only two parental colonies involved, Fig. 10.2A), because the number of eggs available per colony limited the number of crosses that could be performed, while still producing sufficient numbers of offspring for longer-term rearing and outplant. To perform single pairwise crosses, parental colonies needed to be collected and isolated in tanks before spawning, with all spawning and fertilisation work carried out *ex situ* in relatively controlled laboratory conditions (Fig. 10.3B). Done at the scale of this study, it was feasible to do this with a relatively small team of people. In some cases, this approach might not be possible, if for example, spawn collection has to be done *in situ* (Fig. 10.3A) or if natal reefs are far from facilities where spawning and early rearing will take place. The total number of pairwise crosses that can be achieved during a spawning event will depend on the extent of synchronous spawning among colonies and the workforce available for the collection, separation and washing of gametes during their viable time period (<2 hours, Oliver and Babcock, 1992; Omori et al., 2001). In addition, in order to maximise the number of reciprocal pairwise crosses produced requires that each cross is reared independently, and therefore the number of culture vessels can quickly escalate. This then requires increased workforce effort to manage the husbandry during the early post spawning period.

For studies where heritability of the trait has been reported with both sperm and eggs (Liew et al., 2020), then mass crosses (more than two selected parental colonies, Fig. 10.2B) may be a better approach than carrying out many individual crosses. These can be done either by a) pooling gametes of all colonies to produce one culture that is divided into replicate larval rearing tanks, or by b) pooling the sperm of several colonies to fertilize the eggs of a single colony. An advantage of pooling gametes of all colonies is that corals can be allowed to spawn together *en masse* in a single tank reducing the work involved in collecting and separating bundles from many individual colonies. However, culture viability can be compromised by a modest percentage of unfertilized eggs that could originate from a single colony (Pollock et al., 2017). Alternatively, pooling the sperm of several colonies to fertilize eggs from one colony requires collection, separation and washing of gametes as previously described for single pairwise crosses. The benefits of this procedure are that it can be replicated with eggs from all donor colonies, resulting in several distinctive crosses with potentially higher fertilization success, as the concentration of gametes can be better controlled. However, whilst logistical constraints during spawning and fertilisation periods are eased through either colony or sperm pooling, a significant drawback to these approaches is that resulting cohorts produced are a mix of either half or nonrelated kin. Long term post-settlement survival in these kin groups may be lower, compared to fully related offspring, due to negative allogenic interactions following developmental onset of the corals’ immune response (Puill-Stephan et al., 2012). Therefore, trade-offs occur between increased work and facilities required to produce large numbers of pairwise crosses versus simplified logistics and reduced larval rearing costs but potentially increased post settlement mortality.

Scientists have successfully conducted CLP for three decades, and great progress has been achieved in controlling steps associated with the early stages, i.e., controlling spawning times (Craggs et al., 2017), fertilizing gametes *en masse*, and rearing larvae to obtaining coral colonies (Guest et al., 2010; Randall et al., 2020). Thus, one of the more challenging steps in CLP is to settle larvae efficiently onto SUs and enhance their survivorship and growth until the outplanting phase (Fig. 10.4). Our results show an average of four corals per SU 10 days after settlement, resulted in at least one colony after four months *ex situ* husbandry rearing. Despite coral post-settlement survivorship being influenced by settlement density and husbandry conditions (Conlan et al., 2017; Cameron and Harrison, 2020), our results provide a useful guideline for optimal settlement densities when using SUs for CLP. For SUs outplanted at different ages, similar analyses are needed to provide information on the minimum number of larvae to settle to obtain one colony per SU at the age of outplant. Furthermore, restoration initiatives will benefit from knowing the optimal number of larvae per SU that will maximise the probability of obtaining a colony that attains sexual maturity, a factor that will be dependent on the overall survivorship of the outplant. An optimal use of available competent larvae for settlement will be a defining factor of the efficiency of the effort. However, the production of larvae is one of the steps which incurs the lowest costs in the framework (Table 3, SM. 12) so efficiency and cost-efficiency may not go hand in hand. Settling more larvae per SU than needed wastes larvae, whereas, not settling too few larvae per SU lowers yield and increase costs. Evidence thus far indicates that low larval settlement densities can compromise the production of SUs with at least one coral colony at the time of outplanting due to high post-settlement mortality, while high settlement densities can compromise survivorship due to density dependent effects (Doropoulos et al., 2017; Cameron and Harrison, 2020). Additionally, high larval densities promote the formation of chimeras, a factor that needs further research as their implications on colony growth and survivorship are still understudied (however see Rinkevich, 2019).

Husbandry of corals during the early post-settlement stages is a key step for successfully conducting CLP as it has a significant impact on growth and survivorship of corals. Techniques for rearing settlers until they can be outplanted to the reef in sufficiently large numbers remain largely experimental. Early survivorship after settlement is the primary bottleneck in CLP for improving cost-effectiveness. Our results show that extending *ex situ* nursery times increased both survivorship and growth rates of outplants and reduced the costs of the production of coral colonies. Husbandry during the early months has been proven to increase coral survivorship (Baria et al., 2010; Guest et al., 2014; Craggs et al., 2019) and growth (this study) by reducing predation and competition pressures (Doropoulos et al., 2016; Gallagher and Doropoulos, 2016). Moreover, *ex situ* rearing can enhance colony growth by controlling light, water and food quality, and by co-culturing corals with herbivores to limit algal overgrowth (Craggs et al., 2019) and reduce costs (Fig. 10.5A, Table 3, SM. 12 and 13). However, there may also be drawbacks associated with longer nursery times such as: 1) acquisition of a symbiont community that differs from the one naturally occurring at the outplant site (Baums et al., 2019), 2) potential fitness consequences of plastic and epigenetic changes due to exposure to different environmental conditions during nursery rearing, (Parkinson and Baums, 2014), and 3) development of pathogens or diseases that might represent a risk (Sheridan et al., 2013).

Many of the disadvantages encountered in *ex situ* nurseries (see above and van Woesik et al., 2020) can be avoided under *in situ* conditions (Fig. 10.5B), especially if the location of the nursery is close to the outplanting reef. However, the costs associated with *in situ* husbandry are considerably higher than *ex situ* conditions (SM. 4), given the logistic and practical constraints associated with working underwater. The location of the *in situ* nursery in a place with similar environmental conditions to the outplanting site, in a shallow area with good water quality that is protected from storms, and that is of easy access can enhance the survivorship and growth of the corals. Although suitable locations for nurseries may be difficult to find, local knowledge and historical environmental data will improve the chances of locating appropriate sites. One major limitation of *in situ* nurseries is that environmental stressors (i.e. temperature fluctuations, storms) and human activities (e.g. diver or anchor impacts) cannot be completely controlled and may damage the nursery and compromise survivorship of the corals. Further, nursery maintenance and monitoring of coral health is an expensive activity and will be limited by the ease of access to the site. Moreover, enhanced growth rates at *in situ* nurseries can be associated with reductions in skeletal densities (Pratchett et al., 2015), a potential disadvantage for colonies’ performance at the natal reef. These factors highlight the need for developing husbandry-outplanting techniques that reduce husbandry time and increase outplant survivorship. A major improvement is required to reduce these costs in order to successfully scale up SB for assisting reefs to be more resistant to climate change disturbances.

The outplanting phase was the most expensive activity of this framework, highlighting the need for technical development (Fig. 10.6) if restoration is to be scaled up. The method used for outplanting corals in this study is not suitable for larger scale restoration and should not be considered as such as it is laborious, slow, and costly and only appropriate at a scale of tens to hundreds of square metres (Guest et al., 2014). The outplanting methodology should minimize the use of tools and attachment materials (i.e., Coralclip®; Suggett et al., 2019). SUs should be designed so that they are easy to transport, can be rapidly and easy deployed, can be readily attached to the reef substrate (i.e., tetrapods; Chamberland et al., 2017) and, if possible be made of sustainable and ecologically friendly constituents (i.e., sustainable cement; de Brito and Kurda, 2021). The design of SU should enhance coral survivorship until they reach an escape size (i.e., with micro-refugia to protect colonies from grazing pressure) and improve the success of early outplant, reducing husbandry time. Interdisciplinary teams of ecologist, aquarists and engineers are key to developing novel designs of SUs that can be easily deployed in different reef environments (i.e., reef crest, reef flat), which is a critical factor for CLP and SB techniques to be adopted as a management strategy of coral reefs.

The cost analysis reveals that extending husbandry time has a major impact on the efficiency of the propagation effort and reducing the cost of 2.5-year-old colonies five times as a result of early survivorship increases. Total costs could be reduced considerably if post-outplant survivorship is increased at younger ages, thus decreasing husbandry times, hence innovation in this area is fundamental to reduce total costs. Likewise, several strategies could be adopted to reduce costs associated with monitoring, i.e., a) surveying only a representative subset of outplants, b) developing a citizen science program which incorporates the local and tourist community during the monitoring phase (Sinclair et al., 2021), and c) use of innovative technologies like photogrammetry to monitor large reef areas (∼250 m) while minimising time underwater (Lechene et al., 2019; however analysis of the costs of this methodology will need to be computed to compare it with the methodology used in this study). Cost breakdown of CLP is not straightforward, owing to the several factors that vary between coral species, sites, facilities and countries. Yet, doing the exercise of estimating such expenses accounting for types of costs (i.e., capital equipment, consumables, labour) provides a quantitative method to identify the steps that need further development to improve overall efficiency and reduce costs. Reef rehabilitation initiatives using CLP under a SB approach will effectively become a management strategy when techniques are proven to be cost-effective. Presently, CLP remains a very expensive practice (Edwards et al., 2010; Nakamura et al., 2011; Guest et al., 2014) that if implemented can take away funds that could be used for environmental protection and conservation. Hence, coastal managers need to consider carefully if active rehabilitation techniques, such as those presented here, are a prudent use of limited funds. Here we present a framework that combines coral sexual propagation with selective breeding that could be employed as part of initiatives to promote adaptation and resilience of reefs in the Anthropocene.

