## Supplementary tables and figures for "A framework for selectively breeding corals for assisted evolution"

**Supplementary material 1:** Temperature profile during the 7-day heat stress assay performed on fragments of 34 *Acropora digitifera* coral colonies. Two temperature levels were used: a) ambient seawater temperature conditions (tank 1, 2, and 8), and b) heat stress conditions (tank 3, 4, 5, 6 and 7), where temperature increments were performed over a course of 3 days (+2°C on day 1, and +1.5°C on day 3), reaching a daily average temperature of 32.95°C (±0.37) during days 4 to 7.

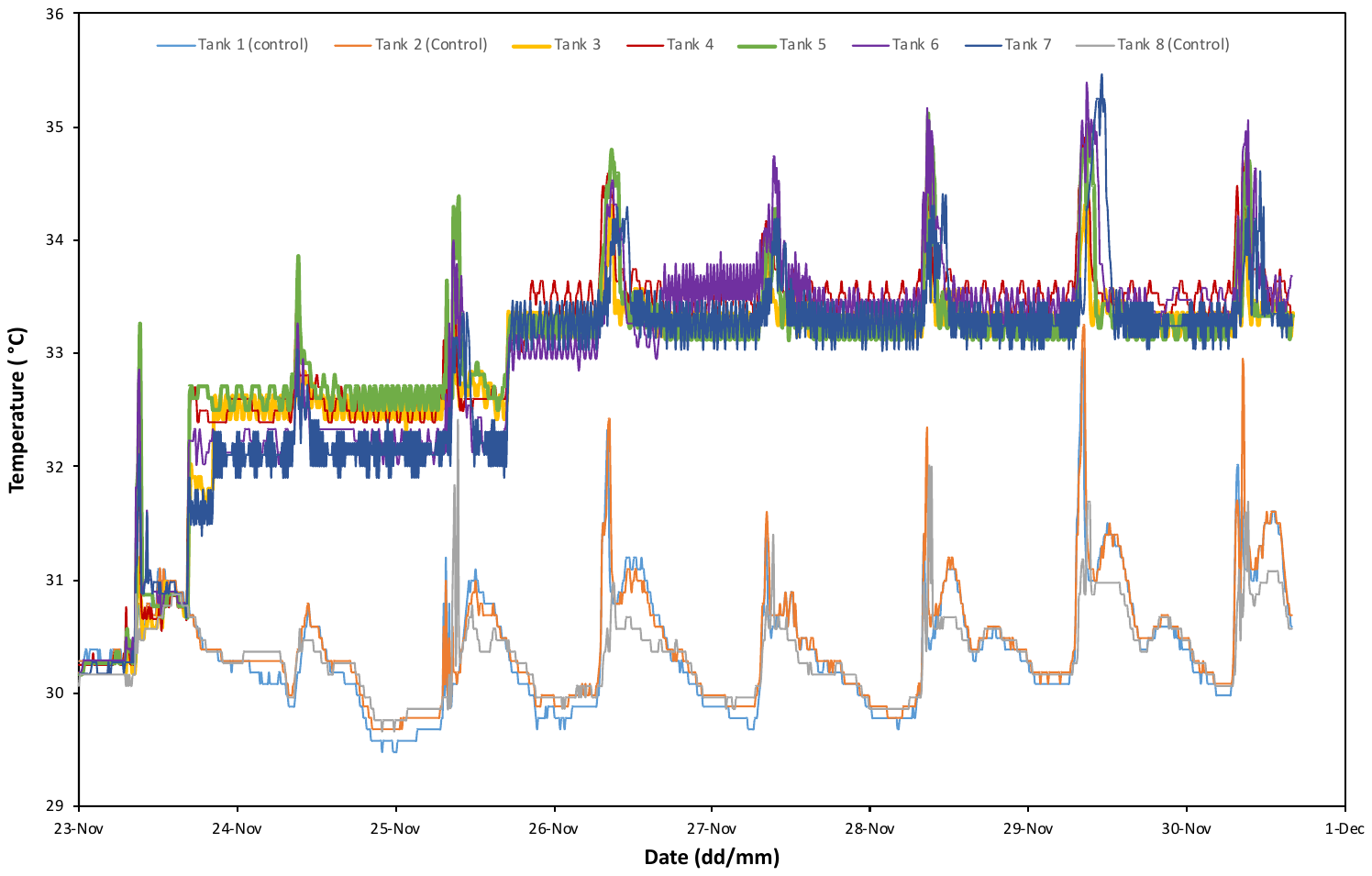

**Supplementary material 2:** Reproductive (gravid: containing mature eggs, or non-gravid: empty polyps) and health (alive, partial mortality, dead or not found) statuses of the 99 tagged colonies one to two days before the full moon in March 2018. Relative heat tolerance (low or high) of the 34 colonies tested in the 7-day heat stress assay (23/11/2017-01/12/2017) is also indicated.

| Relative Heat Tolerance | Date sampled | Reproductive Status | Health status |
| --- | --- | --- | --- |
| High | 29/3/18 | Mature | Alive |
| High | 29/3/18 | Empty | Alive |
| High | 29/3/18 | Empty | Alive |
| High | 29/3/18 | Mature | Alive |
| High | 29/3/18 | Mature | Alive |
| High | 29/3/18 | Empty | Alive |
| High | 29/3/18 | Empty | Alive |
| High | 29/3/18 | Empty | Alive |
| High | 29/3/18 | Mature | Alive |
| High | 29/3/18 | Mature | Alive |
| High | 29/3/18 | Empty | Partial mortality |
| High | 29/3/18 | Mature | Alive |
| Low | 29/3/18 | Mature | Alive |
| Low | 29/3/18 | Mature | Alive |
| Low | 29/3/18 | Empty | Alive |
| Low | 29/3/18 | Mature | Alive |
| Unclassified | 29/3/18 | NA | Dead |
| Unclassified | 29/3/18 | Mature | Alive |
| Unclassified | 29/3/18 | Mature | Alive |
| Unclassified | 29/3/18 | Mature | Alive |
| Unclassified | 29/3/18 | Empty | Alive |
| Unclassified | 29/3/18 | Mature | Alive |
| Unclassified | 29/3/18 | Mature | Alive |
| Unclassified | 30/3/18 | Mature | Alive |
| Unclassified | 29/3/18 | Mature | Alive |
| Unclassified | 30/3/18 | Mature | Alive |
| Unclassified | 30/3/18 | Mature | Alive |
| Unresolved | 29/3/18 | Empty | Alive |
| Unresolved | 29/3/18 | Empty | Alive |
| Unresolved | 29/3/18 | Mature | Alive |
| Unresolved | 29/3/18 | Empty | Alive |
| Unresolved | 29/3/18 | NA | Dead |
| Unresolved | 29/3/18 | Mature | Alive |
| Unresolved | 29/3/18 | Mature | Alive |
| Not tested | 29/3/18 | Mature | Alive |
| Not tested | 29/3/18 | NA | Dead |
| Not tested | 29/3/18 | Empty | Alive |
| Not tested | 29/3/18 | Mature | Alive |
| Not tested | 29/3/18 | Empty | Alive |
| Not tested | 29/3/18 | Empty | Alive |
| Not tested | 29/3/18 | Empty | Alive |
| Not tested | 29/3/18 | Empty | Alive |
| Not tested | 29/3/18 | Empty | Alive |
| Not tested | 29/3/18 | Empty | Alive |
| Not tested | 29/3/18 | Empty | Alive |
| Not tested | 29/3/18 | NA | Not found |
| Not tested | 29/3/18 | Empty | Alive |
| Not tested | 29/3/18 | Empty | Alive |
| Not tested | 29/3/18 | Empty | Alive |
| Not tested | 29/3/18 | Empty | Alive |
| Not tested | 29/3/18 | Empty | Alive |
| Not tested | 29/3/18 | Empty | Alive |
| Not tested | 29/3/18 | Empty | Alive |
| Not tested | 29/3/18 | Mature | Alive |
| Not tested | 29/3/18 | Mature | Alive |
| Not tested | 29/3/18 | Empty | Alive |
| Not tested | 29/3/18 | Empty | Alive |
| Not tested | 29/3/18 | Empty | Alive |
| Not tested | 29/3/18 | Empty | Alive |
| Not tested | 29/3/18 | NA | Dead |
| Not tested | 29/3/18 | Empty | Alive |
| Not tested | 29/3/18 | Empty | Alive |
| Not tested | 29/3/18 | Empty | Alive |
| Not tested | 29/3/18 | NA | Dead |
| Not tested | 29/3/18 | Mature | Alive |
| Not tested | 29/3/18 | Empty | Alive |
| Not tested | 29/3/18 | Empty | Alive |
| Not tested | 29/3/18 | NA | Dead |
| Not tested | 29/3/18 | Empty | Alive |
| Not tested | 29/3/18 | Empty | Alive |
| Not tested | 29/3/18 | Empty | Alive |
| Not tested | 29/3/18 | Empty | Alive |
| Not tested | 29/3/18 | Empty | Alive |
| Not tested | 29/3/18 | Mature | Alive |
| Not tested | 29/3/18 | Mature | Alive |
| Not tested | 29/3/18 | Mature | Alive |
| Not tested | 29/3/18 | Empty | Alive |
| Not tested | 29/3/18 | Mature | Alive |
| Not tested | 29/3/18 | Empty | Alive |
| Not tested | 29/3/18 | Empty | Alive |
| Not tested | 29/3/18 | Empty | Partial mortality |
| Not tested | 29/3/18 | Empty | Alive |
| Not tested | 29/3/18 | Mature | Alive |
| Not tested | 29/3/18 | NA | Dead |
| Not tested | 29/3/18 | Mature | Alive |
| Not tested | 29/3/18 | Empty | Partial mortality |
| Not tested | 29/3/18 | Empty | Partial mortality |
| Not tested | 29/3/18 | Empty | Partial mortality |
| Not tested | 29/3/18 | Empty | Alive |
| Not tested | 30/3/18 | Empty | Alive |
| Not tested | 30/3/18 | Empty | Alive |
| Not tested | 30/3/18 | Mature | Alive |
| Not tested | 30/3/18 | Empty | Alive |
| Not tested | 29/3/18 | Mature | Alive |
| Not tested | 30/3/18 | Mature | Alive |
| Not tested | 30/3/18 | Mature | Alive |
| Not tested | 30/3/18 | Mature | Alive |
| Not tested | 29/3/18 | NA | Dead |
| Not tested | 30/3/18 | Mature | Partial mortality |

**Supplementary material 3:** Upper view lay out of the setup for the cones used for larvae rearing.

**Supplementary material 4:**

Labour was divided into two skill levels, scientific advisor and trained manual labour at an annual salary of US$38,000 (Postdoctoral Research Associate salary in the United Kingdom) and US$12,000 (Trained Manual Labour salary in Palau), respectively. Bench fees covering access to a seawater supply (sand filtered), electricity, usage of tanks, and laboratory facilities were not considered as these increase equally the total cost of the evaluated scenarios and will vary considerably among countries, institutions or facilities. Hardware prices are from stores in Palau at the time of the study. Capital equipment that was used for different activities was pro-rated over five years among categories based on person-days usage. The cost analysis was based on the maximum number of eggs that can be reared in 24 larval rearing tanks (4500 larvae per tank = 108,000 eggs). The number of colonies needed for producing 108,000 eggs was estimated from the literature for *Acropora* sp. (SM. 5). The number of people needed during the spawning night, outplanting and monitoring was four (two scientific advisors and two trained manual labours) to increase the success and efficiency of those activities and decrease the total costs. Fertilization success was set to 95% (Humanes et al. 2016) survival to competency 90% (Humanes et al. 2016), and the percentage of larvae that settle on the Substrate Units (SUs) was 50% (Cameron and Harrison 2020). The number of larvae offered per SU was 25, a rather conservative value considering that our density settlement analysis suggest that only four settlers per SU are needed to obtain at least one juvenile coral after four months of *ex situ* rearing. Survivorship under *ex situ* rearing conditions until outplanting was 90% and 78% for each outplant age (five and 11 months respectively), values estimated during the 2020 spawning event in pre-conditioned tanks with the addition of juvenile trochus snails and feeding three times a week. Outplanting was considered to occur in the natal reef of this study (depth between 0 and 4 m) allowing two divers to outplant 50 SUs per dive (25 SU outplanted per 10 m long transects). Monitoring was costed twice for each outplant and photographs from 200 colonies were stipulated during each monitoring to estimate growth.

**Supplementary material 5:** Number of eggs produced per colony by Acropora species in different studies.

| **Reference** | **Species** | **Colonies collected** | **total #eggs released** | **#eggs released per colony** |
| --- | --- | --- | --- | --- |
| Baria‐Rodriguez et al. (2019) | *Acropora granulosa* | 19 | 1,000,000 | 52,632 |
| Guest et al. (2014) | *Acropora millepora* | 3 | 120,000 | 40,000 |
| Ligson et al. (2019) | *Acropora verweyi* | 11 | 300,000 | 27,273 |
| Nakamura et al. (2011) | *Acropora tenuis* | 14 | 237,000 | 16,929 |
| Maximum |  |  |  | 52,632 |
| Minimum |  |  |  | 16,929 |

**Supplementary material 6:** Reproductive (gravid: containing mature eggs, or non-gravid: polyps empty) statuses of the 9 colonies assessed on the 31 of March 2020, 6 days before the spawning event.

| Colony | Status |
| --- | --- |
| G | Empty |
| H | Empty |
| I | Empty |
| J | Empty |
| K | Empty |
| L | Empty |
| M | Empty |
| N | Empty |
| O | Empty |

**Supplementary material 7:** Number of substrate units (SUs) present in each of the eight settlement tanks with stocked at three levels of larval densities (10, 25 and 50 larvae per SU).

| **Larval density** | **Settlement tank replicate** | **Number of substrate units (SUs)** |
| --- | --- | --- |
| **10** | 1 | 4 |
|  | 2 | 10 |
|  | 3 | 9 |
|  | 4 | 8 |
|  | 5 | 9 |
|  | 6 | 10 |
|  | 7 | 8 |
|  | 8 | 7 |
| **25** | 1 | 8 |
|  | 2 | 10 |
|  | 3 | 10 |
|  | 4 | 10 |
|  | 5 | 6 |
|  | 6 | 9 |
|  | 7 | 7 |
|  | 8 | 6 |
| **50** | 1 | 9 |
|  | 2 | 9 |
|  | 3 | 10 |
|  | 4 | 7 |
|  | 5 | 10 |
|  | 6 | 6 |
|  | 7 | 9 |
|  | 8 | 9 |

**Supplementary material 8:** Model validation test results show the ability of GLMMs to represent zero-inflated response data for two models: 1) SD ~ LCD, and 2) C4D ~ SDnz. These models are written in the “Response ~ Predictor + Random Tank Effect” format (see tables below), where LCD is larval culture density, SD is settlement density, C4D is four-month coral density, and SDnz is non-zero settlement density. Models were fit using either Poisson or negative binomial variance structures. The true percentage of zeros from measured data (red diamond) are compared to simulation histograms, showing the distribution of percentages of zeros from 1000 simulated response variables for each of the four models. True values lie the 95% confidence intervals of simulated values (dashed red lines) for all Poisson models, while negative binomial models all performed better, also shown by the dispersion statistic (D) with an optimal value of one.

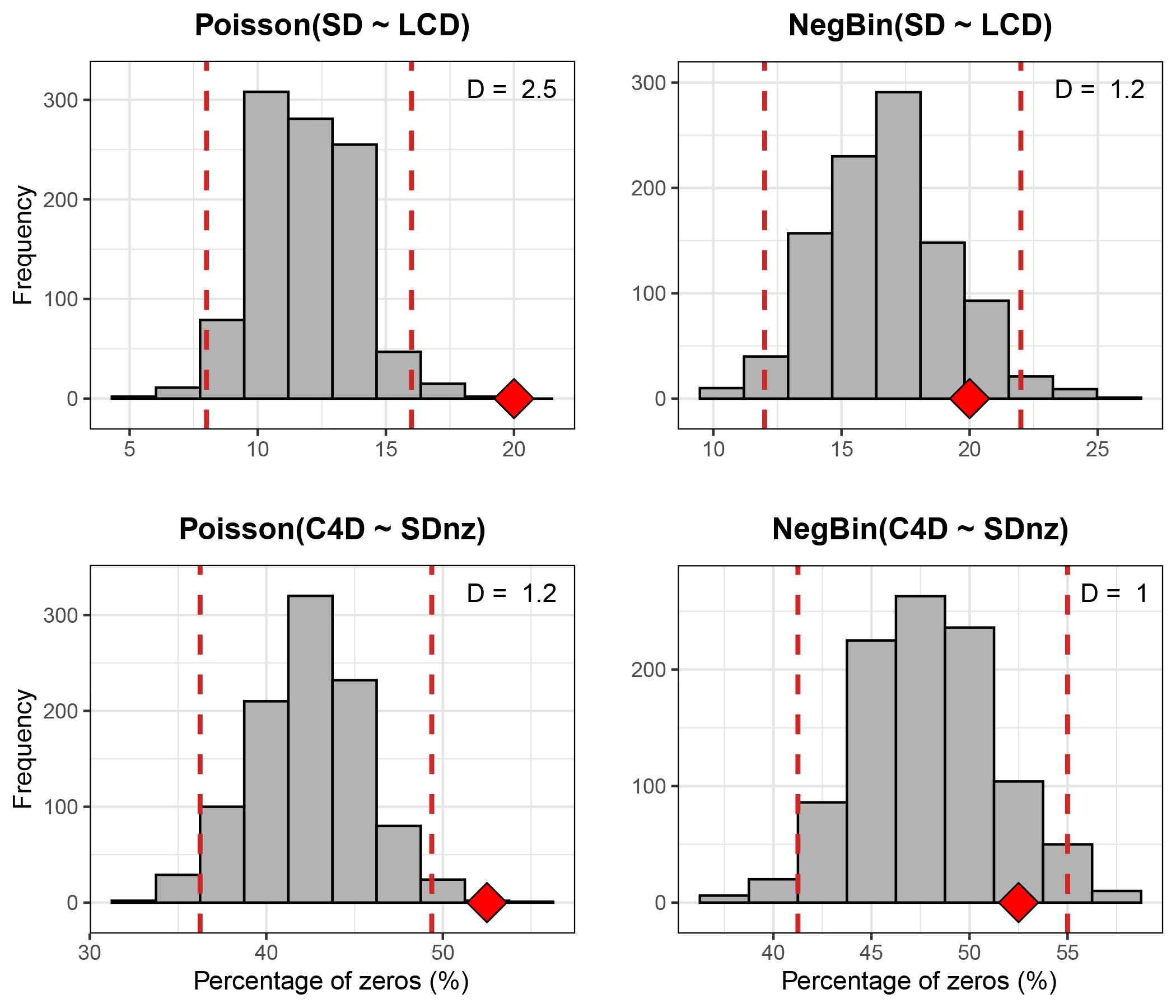

**Supplementary material 9:** Results for *Acropora digitifera* showing the relationship between larval density (3-levels) on 10-days old settler density per substrate unit (SU) based on generalized mixed models (GLMM) with the log-negative binomial function with settlement tank as random factor. Significance is shown at p<0.05 in bold. Model: Number of settlers density per SU ~ larval culture density + (1|settlement tank), output on logarithmic scale.

|  | Estimate | Std. Error | z value | p |
| --- | --- | --- | --- | --- |
| Intercept | 0.52 | 0.12 | 4.16 | **<0.001** |
| 25 larvae per SU | 0.45 | 0.17 | 2.73 | **<0.01** |
| 50 larvae per SU | 1.1 | 0.16 | 7.07 | **<0.001** |

**Supplementary material 10:** Results for *Acropora digitifera* showing relationship between 10-days old settler density per SU (non-zero) on four-months old coral density per substrate unit (SU) based on generalized mixed models (GLMM) with the log-negative binomial function with *ex situ* nursery tank as random factor. Significance is shown at p<0.05 in bold. Model: Number of corals after four months per SU ~ log(number of settlers per SU) + (1|ex situ nursery tank), output on logarithmic scale.

|  | Estimate | Std. Error | z value | p |
| --- | --- | --- | --- | --- |
| Intercept | -1.44 | 0.31 | -4.63 | **<0.001** |
| log(number of settlers per SU) | 1.02 | 0.14 | 7.12 | **<0.001** |

**Supplementary material 11:** Results for *Acropora digitifera* showing how planar area differs depending on rearing strategy (outplanted to the reef at five and 11-month old or moved to the *in situ* nursery at 13-months old) based on general mixed models (GLMM) with the log-gamma function, with outplant method as fixed factor and cross as random factor. Significance is shown at p<0.05 in bold. Model: Planar area ~ rearing strategy + (1|cross).

|  | Estimate | Std. Error | t value | P |
| --- | --- | --- | --- | --- |
| Intercept | 0.16 | 0.22 | 0.72 | 0.47 |
| 11-month old outplant to the reef | 1.01 | 0.24 | 4.16 | **<0.001** |
| Transferred to the *in situ* nursery | 2.25 | 0.26 | 8.76 | **<0.001** |

**Supplementary material 12:** Cost of sexually propagating *Acropora digitifera* corals using a selective breeding framework with corals reared under *ex situ* nursery tanks until five months old when they were outplanted to the reef. The capital equipment is pro-rated over five years.

| **Category** | **Capital equipment** | **Consumables** | **Labour** | **Total** | **% total cost** |
| --- | --- | --- | --- | --- | --- |
| Coral collection | $413 | $785 | $312 | $1,510 | 6 |
| Spawning to competency | $1,353 | $803 | $1,274 | $3,430 | 14 |
| Settlement | $177 | $624 | $213 | $1,013 | 4 |
| Ex situ nursery rearing | $1,117 | $254 | $379 | $1,750 | 7 |
| Outplanting at five months old | $365 | $5,490 | $2,412 | $8,267 | 35 |
| Monitoring and maintenance | $559 | $4,019 | $3,269 | $7,847 | 33 |
| **Grand Total** | $3,984 | $11,975 | $7,857 | $23,817 | **100** |

**Supplementary material 13:** Calculations of sexually propagating *Acropora digitifera* corals using a selective breeding framework with corals reared under *ex situ* nursery tanks until 11-months old when they were outplanted to the reef. The capital equipment is pro-rated over five years (SM. 13 Cost analysis larval rearing methods 2018_11 month outplant.xlsx).

**Supplementary material 14:** Calculations of sexually propagating *Acropora digitifera* corals using a selective breeding framework with corals reared under *ex situ* nursery tanks until five months old when they were outplanted to the reef. The capital equipment is pro-rated over five years (SM. 14 Cost analysis larval rearing methods 2018_5 month outplant.xlsx).

**Supplementary material 15:** Cost of building and maintaining nurseries over a period of 11 months. The capital equipment is pro-rated over five years.

| **Category** | **Capital equipment** | **Consumables** | **Labour** | **Total** | **Cost per SU** |
| --- | --- | --- | --- | --- | --- |
| Ex situ nursery rearing ~2,500 SUs | $1,592 | $254 | $575 | $2,421 | $0.97 |
| In situ nursery rearing for ~1,700 SUs | $384 | $7,413 | $1,977 | $9,774 | $5.75 |

**Supplementary material 16:** Calculations of building and maintaining an *in situ* nursery for 1,700 SUs over a period of 11 months. The capital equipment is pro-rated over five years (SM. 16 Cost analysis larvae rearing methods 2018_In-situ nursery.xlsx).
